# Autism genetics perturb prenatal neurodevelopment through a hierarchy of broadly-expressed and brain-specific genes

**DOI:** 10.1101/2020.05.23.112623

**Authors:** Vahid H. Gazestani, Austin WT Chiang, Eric Courchesne, Nathan E. Lewis

**Author notes:** Correspondence to Nathan E. Lewis and Eric Courchesne.

## Abstract

Numerous genes are associated with autism spectrum disorder (ASD); however, it remains unclear how most ASD risk genes influence neurodevelopment and result in similar traits. Recent genetic models of complex traits suggest non-tissue-specific genes converge on core disease genes; so we analyzed ASD genetics in this context. We found ASD risk genes partition cleanly into broadly-expressed and brain-specific genes. The two groups show sequential roles during neurodevelopment with broadly-expressed genes modulating chromatin remodeling, proliferation, and cell fate, while brain-specific risk genes are involved in neural maturation and synapse functioning. Broadly-expressed risk genes converge onto brain-specific risk genes and core neurodevelopmental genes through regulatory networks including PI3K/AKT, RAS/ERK, and WNT/*β*-catenin signaling pathways. Broadly-expressed and brain-specific risk genes show unique properties, wherein the broadly-expressed risk gene network is expressed prenatally and conserved in non-neuronal cells like microglia. However, the brain-specific gene network expression is limited to excitatory and inhibitory neurons, spanning prenatal to adulthood. Furthermore, the two groups are linked differently to comorbidities associated with ASD. Collectively, we describe here the organization of the genetic architecture of ASD as a hierarchy of broadly-expressed and brain-specific genes that disrupt successive stages of core neurodevelopmental processes.

## INTRODUCTION

Autism spectrum disorder (ASD) is a common and largely heritable neurodevelopmental disorder characterized by deficits in social interaction and repetitive behaviors. Genetic studies have implicated hundreds of ASD risk (rASD) genes (De Rubeis et al., 2014; Iossifov et al., 2014; Krumm et al., 2015; Sanders et al., 2015; Turner et al., 2017; Yuen et al., 2017) with peak expression at prenatal and early postnatal ages, indicating the early onset of the disorder(Courchesne et al., 2019; Grove et al., 2019; Parikshak et al., 2013; Satterstrom et al., 2020; Willsey et al., 2013). However, rASD genes demonstrate astonishing heterogeneity with different neurodevelopmental expression patterns and enrichment in diverse biological processes (de la Torre-Ubieta et al., 2016; Duda et al., 2018; Krishnan et al., 2016; Li et al., 2018; Parikshak et al., 2013; Pinto et al., 2014). The extreme heterogeneity of rASD genes pose a formidable challenge in connecting ASD genetics to the molecular mechanisms leading to dysregulated development and the clinical phenotypes observed in the ASD population.

To decipher the complex genetic structure of ASD, studies have attempted to identify neuronal cell contexts in which rASD genes, as a single group, demonstrate strong (co-)expression (Chang et al., 2015; Krumm et al., 2014; Pinto et al., 2010; Satterstrom et al., 2020; Willsey et al., 2013). These models, in essence, hypothesize that the whole group of heterogeneous rASD genes form a coherent network in specific neuronal cell types and at a defined spatio-temporal space to perform a specific biological function. These single-group rASD network models, therefore, place the perturbation of such a network at the core of ASD pathobiology. Although valuable, such models ignore potential impacts of ASD genetics in other cellular or developmental contexts. For example, rASD genes are linked to biological processes related to non-neuronal cell types including microglia, astrocytes, and leukocytes (Ballas et al., 2009; Derecki et al., 2012; Gupta et al., 2014; Kasah et al., 2018; Pramparo et al., 2015; Suzuki et al., 2013; Voineagu et al., 2011). Likewise, genetic variants related to brain processes, diseases, and disorders can modulate gene expression in cell types other than neurons and tissues other than brain (Lin et al., 2018; Lombardo et al., 2018; Pardinas et al., 2018; Wright et al., 2014).

Recent studies present an alternate model of complex traits where the functional impact of genetic aberrations propagates through gene regulatory networks to converge on downstream core processes related to the trait (Boyle et al., 2017; Califano and Alvarez, 2017; Courchesne et al., 2020; Gazestani et al., 2019; Iakoucheva et al., 2019; Liu et al., 2019). In contrast to the single-group rASD network model, if the polygenic and omnigenic models are relevant to ASD, they would suggest that many risk genes are not necessarily part of underlying neural mechanisms in ASD. Rather many rASD genes would be broadly-expressed across many tissues, but regulate core neurodevelopmental processes underlying ASD (Boyle et al., 2017; Califano and Alvarez, 2017; Courchesne et al., 2020; Gazestani et al., 2019). Moreover, by emphasizing the gene regulatory networks, these alternative models suggest that, to the extent that the gene regulatory networks are preserved, the dysregulation could also be reflected in non-neuronal cells (Gazestani et al., 2019). Supporting these ideas, many rASD genes are broadly-expressed across tissues and strongly enriched for gene expression regulators such as chromatin remodelers, transcription factors, and modulators of signaling pathways (Courchesne et al., 2020; Courchesne et al., 2019; Werling et al., 2020). Similarly, genome wide association studies (GWAS) and whole genome sequencing both highlight the important role of broadly functional eQTLs and gene regulatory networks in ASD liability (Brandler et al., 2018; Courchesne et al., 2020; Grove et al., 2019).

The fundamental unanswered question, therefore, is how heterogenous rASD genes connect with the molecular, cellular, and developmental perturbations in ASD. Is there a unifying model of the ASD genetic architecture, or do different groups of rASD genes connect to different molecular and developmental perturbations? We hypothesized that examining the connection of broadly-expressed and brain-specific rASD genes to molecular perturbations in ASD neuron models would elucidate how the heterogeneous genetics converge to form similar traits, and highlight how the genetic architecture of ASD is organized. We investigated the importance of gene regulatory networks in connecting ASD genetics with the underpinning molecular perturbations and the pathobiology of ASD. We demonstrate there are two distinct classes of rASD genes: brain-specific and broadly-expressed genes. These two classes of genes show distinct temporal expression patterns in the developing brain. They also form two separate gene networks, but the networks are connected by signaling pathways that transmit perturbations in broadly-expressed genes to the core neurodevelopmental processes underlying ASD that include brain-specific risk genes, ultimately leading to dysregulation of neurodevelopment and emergence of ASD characteristics. Thus, we present a comprehensive and unified model describing the genetic architecture of ASD.

## RESULTS

### ASD risk genes show two distinct expression patterns: broadly-expressed and brain-specific

There are many brain-specific rASD genes, but many leading risk genes are transcriptional regulators expressed ubiquitously across tissues (e.g. CHD8); thus, we systematically tested how rASD genes are expressed across tissues. We focused on rASD genes relevant to ASD pathobiology that were identified by at least two datasets according to SFARI database (Abrahams et al., 2013). To systematically classify rASD genes as brain-specific or broadly-expressed across tissues, we evaluated the expression patterns of rASD genes across 26 adult tissues from 10,259 GTEx RNA-Seq samples (Consortium, 2015). 83% (193 out of 232) of rASD genes were expressed in cortex samples (Fig 1A). In comparison, only 59% (344 out of 584) of mutated genes in healthy siblings of ASD subjects (ASDsib genes) were expressed in the same cortex samples (OR: 3.4; *P*: 1.2×10^−11^; Fig S1-S2).

**Fig 1.**
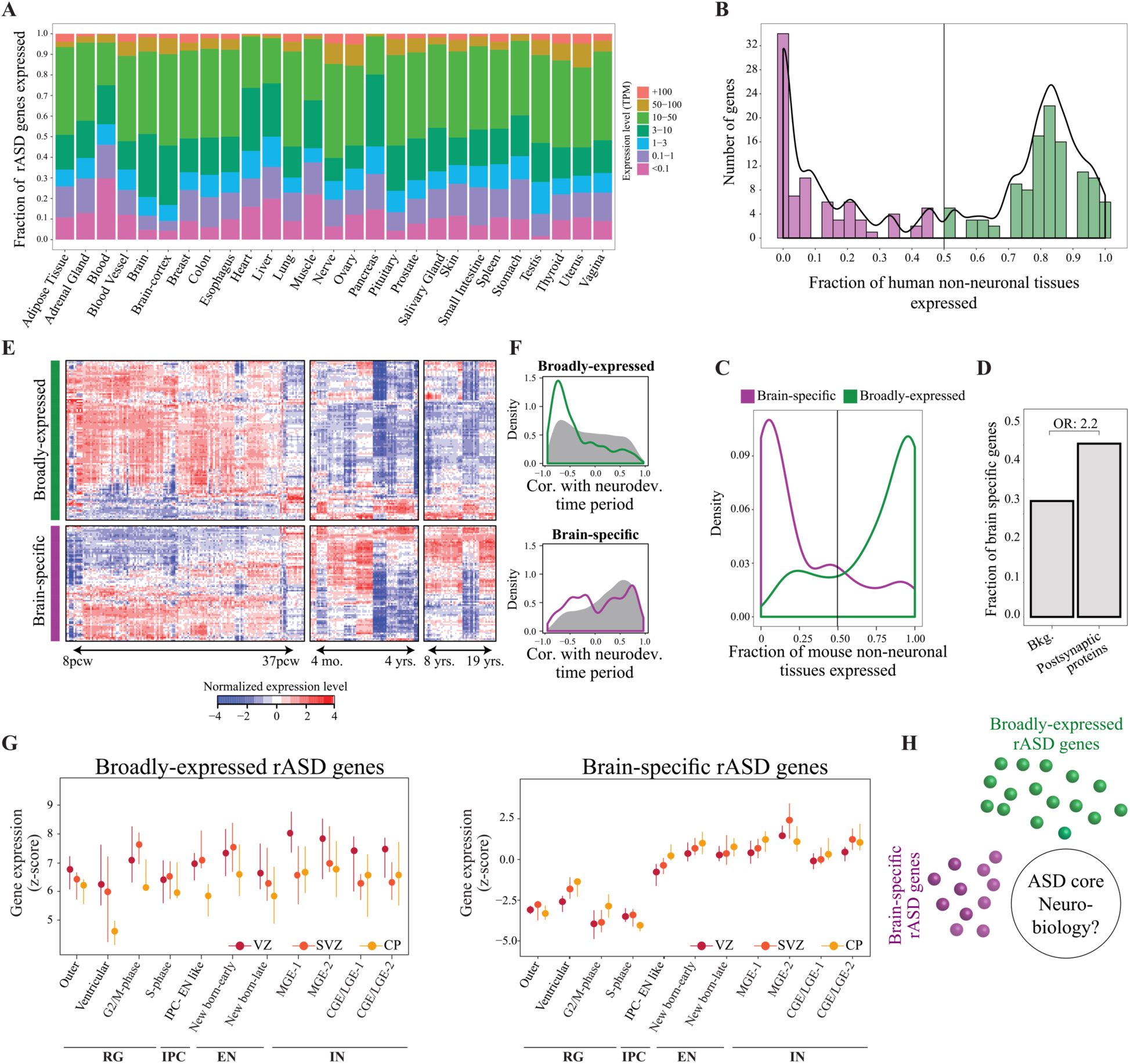
ASD risk genes include both broadly-expressed and brain-specific genes with temporally different expression patterns. **A)** Gene expression level of rASD genes across human tissues. Most rASD genes are strongly expressed in the brain, and many are broadly expressed across other tissues. **B)** rASD genes include broadly-expressed and brain-specific genes. For the rASD genes expressed in the adult cortex, we calculated the fraction of non-neuronal tissues wherein the genes are expressed. **C)** Homologs to human broadly-expressed rASD genes are widely expressed across mouse tissues, while brain-specific gene homologs exhibit limited expression across non-neuronal mouse tissues. **D)** Genes involved in synapse structure and function are enriched for brain-specific genes, indicating the functional relevance of the gene categorization based on the tissue expression patterns. **E)** Neocortex gene expression data for broadly-expressed and brain-specific rASD genes spanning prenatal and postnatal ages. While broadly-expressed genes are gradually downregulated during the neurodevelopment, many brain-specific genes show strong expression in late prenatal and postnatal ages. **F)** The neocortex expression patterns of broadly-expressed and brain-specific rASD genes are compared to all broadly-expressed and brain-specific genes in general (grey). The correlation of gene expression patterns with the prenatal and early postnatal time periods (8 pcw to 2 years after birth) were calculated using the biweight midcorrelation efficiently metric. **G)** Single cell data illuminate the different expression trajectories of broadlyexpressed and brain-specific rASD genes at prenatal time periods. While the expression of broadly-expressed rASD genes are mostly decreased during laminae transitions of cells across cortex, the expression of brain-specific rASD genes are strongly affected by cell type differentiation patterns. The expression of the brain-specific genes is also affected by the laminae transitions of the cells, but in an opposite direction to the broadly-expressed rASD genes. Circles represent the median z-score from GSEA analysis, and lines illustrate associated interquartile ranges. **H)** rASD genes are composed of two groups of broadly-expressed and brain-specific genes. We find the two groups are regulated by different transcriptional programs. However, the question remains if and how the genes in two groups are connected to each other and to the core neurobiology of ASD.

We subsequently quantified tissue specificity of the 193 expressed rASD genes by calculating the fraction of non-brain tissues expressing each rASD gene. We found a clear bimodal distribution with the peaks at zero and 0.85 (0 = only in brain; 1 = in all tissues), suggesting two major groups of rASD genes (Fig 1B). We divided the 193 rASD genes into two groups –broadly-expressed and brain-specific– with broadly-expressed genes expressed in ≥50% of non-brain tissues (Boyle et al., 2017; Courchesne et al., 2020). We found 86% of brain-specific genes are expressed in ≤25% of other tissues, suggesting their more exclusive expression in the brain. The relevance of brain-specific genes to brain processes, and neurons more specifically, was further supported by their limited expression in non-brain tissues of mice (Pervouchine et al., 2015), their enrichment for the synapse proteins (OR: 2.2; *P*: 5.3×10^−43^) (Bayes et al., 2011), and their evolutionary conservation patterns across eukaryotes (Fig 1C-D and S1). Importantly, while rASD genes were significantly enriched for the brain-specific genes (OR: 1.4; *P*: 0.017), many rASD genes (112 out of 193 genes) were broadly-expressed.

### Broadly-expressed express earlier in brain development than brain-specific rASD genes

The striking differences in tissue expression of rASD genes raises an important question regarding if they impact different developmental stages in the brain. Thus, we investigated the rASD gene expression patterns during neurodevelopment using the BrainSpan Atlas (BrainSpan, 2016; Kang et al., 2011). In the developing brain, most broadly-expressed rASD genes show greater peak expression in the first and the second trimester followed by substantial downregulation thereafter, compared to all broadly-expressed genes (Fig 1E-F; *P*: 1.30×10^−8^; Wilcoxon-Mann-Whitney test). In contrast, brain-specific rASD genes demonstrate a bimodal gene expression pattern with 47% and 53% of brain-specific rASD genes showing peak expression in “early” prenatal and “late” prenatal and postnatal neurodevelopment, respectively. This bimodal pattern was also different from all brain-specific genes (Fig 1E-F; *P*: 0.035 Wilcoxon-Mann-Whitney test). As further support, we found similar gene expression trajectories for broadly-expressed and brain-specific rASD genes during *in vitro* differentiation of primary human neurons (Fig S3).

We next analyzed the expression of the two rASD gene groups in different cell types at different brain laminae using single cell RNA-Seq data from 10 to 20 post-conception weeks (pcw) (Nowakowski et al., 2017). Broadly-expressed rASD genes were more strongly expressed than the brain-specific rASD genes during this early prenatal period across different cell types, emphasizing their importance in neurodevelopment (Fig S4). Both broadly-expressed and brain-specific rASD genes were more strongly expressed during the early prenatal period than overall broadly-expressed and brain-specific genes, respectively (*P* <2×10^−16^). As the brain develops, neural progenitor cells differentiate into distinct cell types, including inhibitory and excitatory neurons, and start the migration and maturation processes. In this context, broadly-expressed rASD gene expression is primarily regulated by cell transitions in laminae rather than differentiation from one cell type (e.g., neural progenitor) to another (e.g., excitatory neuron, inhibitory neuron) (Fig 1G). Expression of broadly-expressed rASD genes decreased from ventricular zone (VZ) to sub-ventricular zone (SVZ) (*P*: 0.02) and was further reduced in the cortical plate (CP) (*P*: 2.1×10^−7^). In contrast, brain-specific rASD gene expression was most strongly regulated by cell type transitions (Fig 1G). Specifically, brain-specific risk genes showed lowest expression in neural progenitor cells and their expression gradually increased as cells differentiated into mature excitatory and inhibitory neurons. This trend was shared between brain-specific rASD genes that showed both early and late peak expression in the bulk RNA-Seq data from developing brain (Fig S4). In early prenatal brain development, the expression of brain-specific rASD genes was also significantly higher in CP compared with VZ (*P*: 0.001; Fig 1G). The result showing early prenatal expression of broadly-expressed rASD genes and stronger adult expression of brain-specific genes was further replicated in another independent single cell dataset (Li et al., 2018) (Fig S4).

Collectively, our findings suggest broadly-expressed and brain-specific rASD genes are involved in distinct transcriptional programs that change dynamically across different stages of prenatal neural development (Fig 1H).

### Broadly-expressed and brain-specific rASD genes form divergent networks connected through intermediate processes

One hypothesis about how heterogeneous rASD genes result in ASD is that they are highly pleotropic. Consequently, the hypothesis suggests that rASD genes, whether broadly-expressed or brain-specific, form a coherent and interconnected network at specific developmental time points and cell types to govern biological functions relevant to ASD pathobiology. To test this hypothesis, we analyzed broadly-expressed and brain-specific rASD gene networks to evaluate how they connect with one another and neurodevelopmental processes. We constructed context-specific gene networks of the two rASD gene groups using prenatal and early postnatal neocortex RNA-Seq data. Briefly, we obtained physical and regulatory interactions surrounding each rASD gene (Cerami et al., 2011; Chatr-Aryamontri et al., 2017; Fabregat et al., 2016), and constructed “neurodevelopmental networks” by retaining interacting gene pairs with strong co-expression at prenatal and early postnatal ages (Methods). The broadly-expressed and brain-specific rASD gene groups have significantly more common neighbors within their own category than expected by chance, indicating they form dense subnetworks (Fig 2A, S5). We next enumerated gene neighbors shared between the broadly-expressed and brain-specific rASD genes. This analysis demonstrated that the broadly-expressed and brain-specific rASD genes are significantly depleted of common neighbors with each other (Fig 2B), a pattern that was replicated using alternative gene network construction approaches (Fig S5). This suggests that the networks of broadly-expressed and brain-specific rASD genes connect to each other through limited and potentially specific intermediate molecular processes.

**Fig 2.**
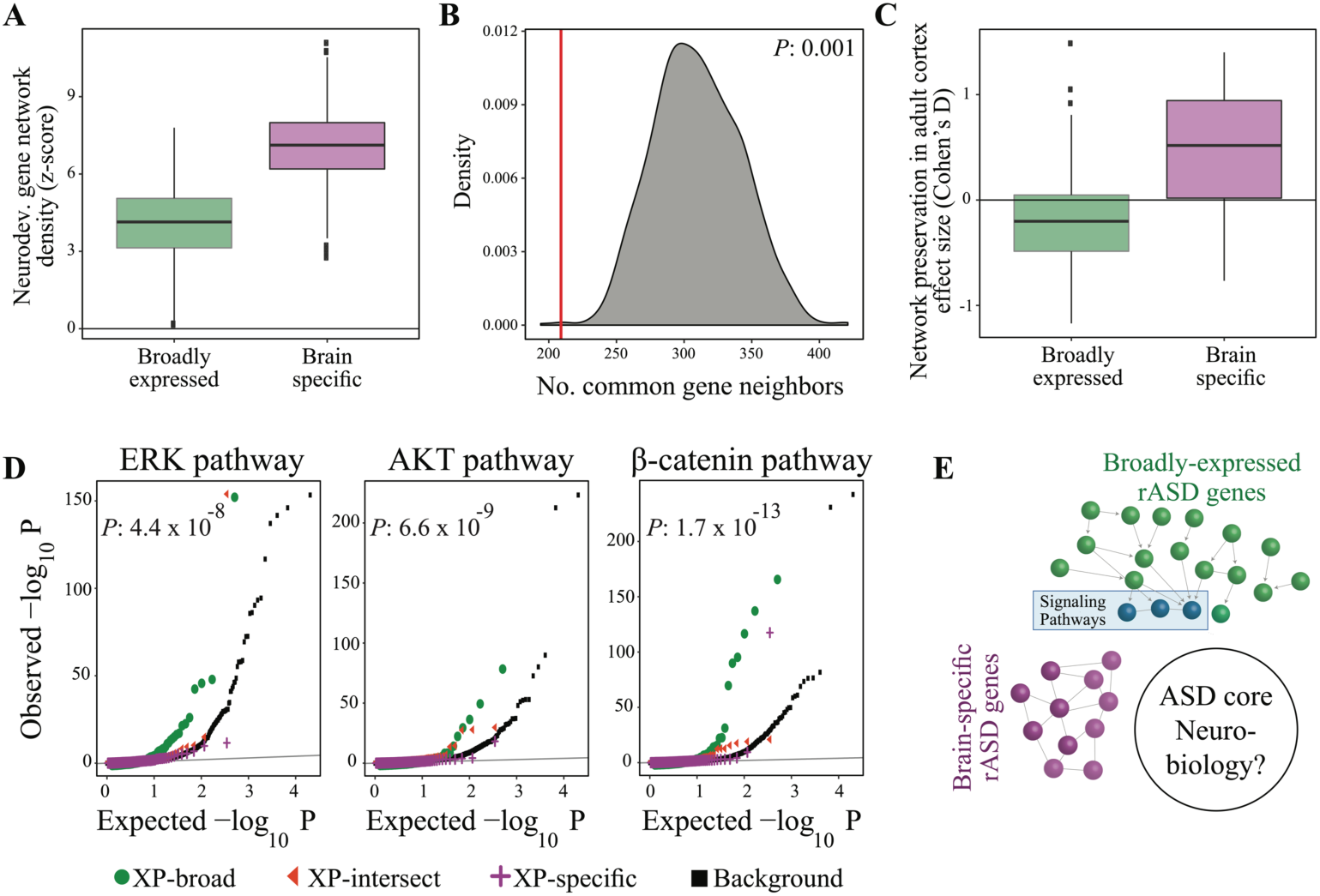
Broadly-expressed and brain-specific rASD genes are involved in different gene networks. **A)** Broadly-expressed and brain-specific rASD genes are involved in dense gene networks. Network density was measured for both groups based on the degree distribution of neighboring genes. If genes in a group share more common gene neighbors, the neighbors would exhibit higher degrees. To test significance, iterating 100 times, degree distributions of neighboring genes were compared to a distribution of randomly selected non-rASD genes from all broadly-expressed or brain-specific genes, respectively. A higher z-score indicates higher within-group network densities. **B)** We enumerated shared gene neighbors between the neurodevelopmental networks of broadly-expressed and brain-specific rASD genes (red line), and found the networks of the two gene sets are significantly separate from each other compared to the background distribution (in gray). Empirical *P* estimated by enumerating shared neighbors from the background distributions. **C)** The neurodevelopmental gene network of brain-specific rASD genes, but not the broadly-expressed genes, is significantly preserved in adult cortex tissue. For each broadly-expressed and brain-specific gene in the neurodevelopmental gene networks, we quantified how much its interactions are preserved between the developing neocortex and adult cortex. A zero effect size indicates that the interactions of a gene in the network of interest are preserved no more than expected by chance and higher effect sizes indicating higher preservation of the interactions. Adult and neurodevelopmental expression data were downloaded from GTEx and BrainSpan, respectively. **D)** The network of broadlyexpressed rASD genes is strongly enriched for the regulators of the PI3K/AKT, RAS/ERK and WNT/*β*-catenin signaling pathways. The comparison between the *P* distribution of XP-broad and the background genes are represented in the panels (Wilcoxon-Mann-Whitney test). **E)** Networks of broadly-expressed and brain-specific genes are separate from each other, though broadly-expressed rASD genes also regulate of PI3K/AKT, RAS/ERK and WNT/*β*-catenin signaling pathways. However, there is still a need to connect these genes to the core neurodevelopmental processes perturbed in ASD.

### Only brain-specific rASD gene networks are continually active in the adult brain

ASD has a prenatal or very early postnatal onset, but some mouse model studies suggest that postnatal treatments targeting synaptic plasticity can ameliorate symptoms and improve brain performance (Delorme et al., 2013; Sahin and Sur, 2015). To see this postnatal responsiveness could be due to a preservation of dysregulated gene networks beyond early development, we tested if rASD genes remain strongly connected to their gene neighbors in the adult cortex. Analysis of GTEx data indicated that the neurodevelopmental network of brain-specific rASD genes is significantly preserved in adult cortex, while the neurodevelopmental network of broadly-expressed rASD genes is not preserved (Fig 2C). Thus, the network of broadly-expressed rASD genes mostly functions during ASD prenatal brain formation. In contrast, the brain-specific rASD gene network remains into later postnatal life. This implies that mutations perturbing the broadly-expressed gene network impact neurodevelopment mostly prenatally, while perturbation of brain-specific gene networks could have an ongoing impact in neuron functions into adulthood.

### Broadly-expressed rASD genes are upstream regulators of ASD signaling pathways and brain-specific rASD genes

We next investigated the biological roles of rASD gene groups. We employed a network propagation approach to identify network neighbors that are strongly (FDR<0.1) connected to broadly-expressed and brain-specific rASD genes. This identified biological processes immediately modulated by rASD genes, involving 355 genes connected to the broadly-expressed and brain-specific rASD genes, collectively. We next categorized the rASD genes and their 355 gene neighbors into three non-overlapping groups of expanded (XP)-broad, XP-specific, and XP-intersect genes. XP-broad and XP-specific networks included all broadly-expressed and brain-specific rASD genes, plus their significantly connected broadly-expressed and brain-specific neighbors, respectively. XP-intersect genes include the neighbors that have connections with the opposing rASD gene groups (i.e., broadly-expressed neighbors that connect to brain-specific rASD genes, and vice versa), and thus represents a bridge between broadly-expressed and brain-specific genes.

XP-broad genes are enriched for processes related to chromatin remodeling, gene expression regulation, cell proliferation, and cell fate determination (Fig 3A). XP-specific genes are enriched for processes related to neurogenesis, neuronal maturation, synapse formation and functioning (Fig 3A). Early-expressing XP-specific genes are enriched in processes related to the neurogenesis, neurite outgrowth, and spine organization, while late-expressing XP-specific genes are mostly involved in synaptic plasticity and synapse functioning (Fig 3A). The XP-intersect network demonstrated PI3K/AKT, RAS/ERK, and WNT/*β*-catenin signaling pathways as key routes that connect XP-broad and XP-specific networks (Fig 3A). This is consistent with previous evidence that these signaling pathways may be perturbed in ASD (Courchesne et al., 2019; Krumm et al., 2014; Sahin and Sur, 2015) and involved in multiple stages of prenatal development from neuronal proliferation to synapse formation and function (Courchesne et al., 2019).

**Fig 3.**
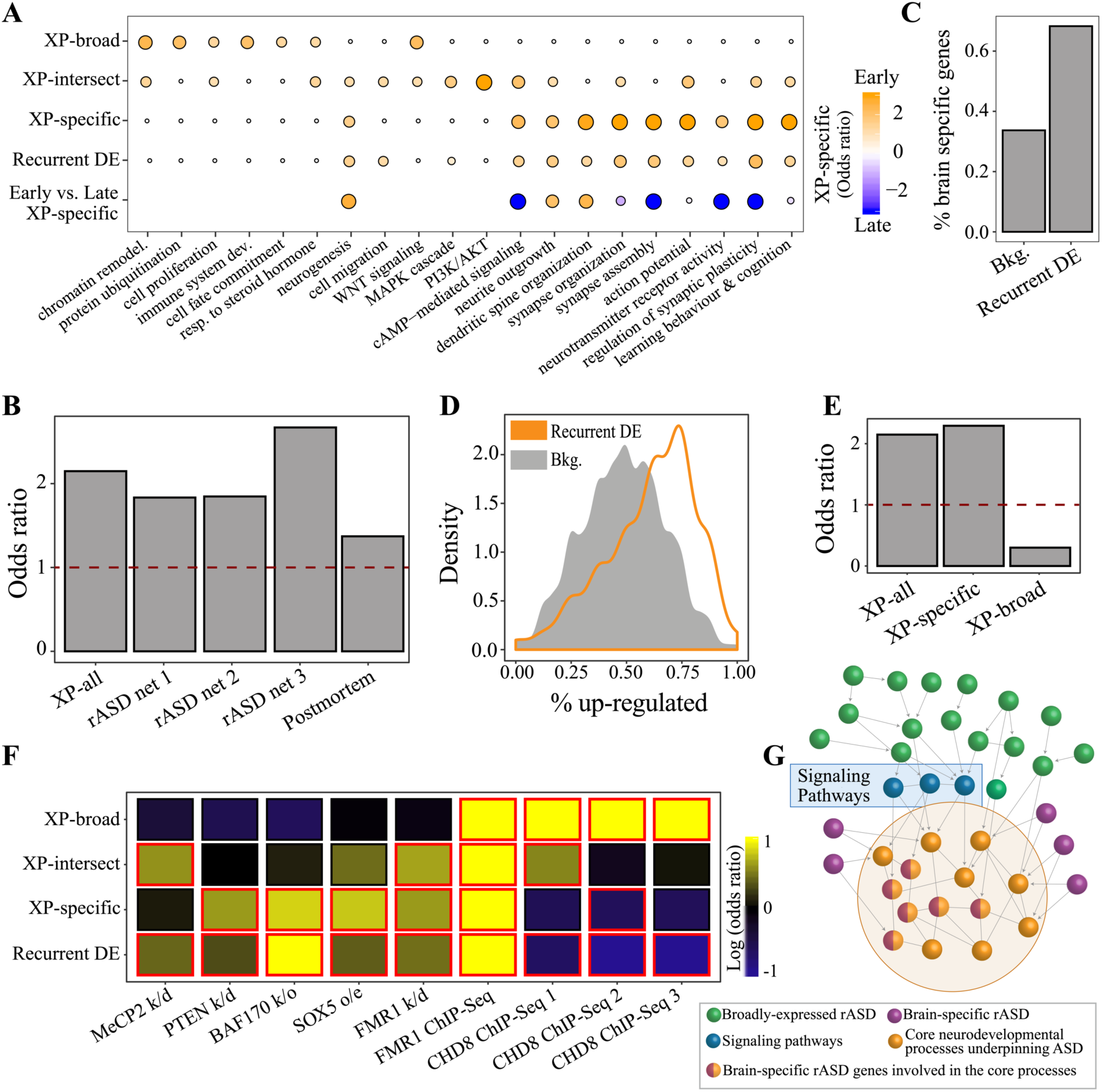
Genes perturbed in ASD overlap with the brain-specific rASD genes. **A)** Gene ontology enrichment analysis of the neurodevelopmental networks of broadly-expressed and brain-specific rASD genes. Broadly-expressed rASD genes are enriched for processes involved in early neural development, while brain-specific gene processes are mostly related to later developmental stages. The genes connecting broadly-expressed and brain-specific networks are enriched in the PI3K/AKT, RAS/ERK, and WNT/*β*-catenin signaling pathways. Processes recurrently differentially expressed in hiPSC-derived neuronal models of ASD overlap substantially with brain-specific rASD gene processes. The last row compares the enrichment in early and late expressing brain-specific rASD genes. Consistent with the neurodevelopmental expression patterns, the early-expressing brain-specific rASD genes are preferentially enriched for neurogenesis, neurite outgrowth, and dendrite spine development (Yellow), while the late brain-specific genes are enriched synaptic plasticity and function (Blue). **B)** Recurrent DE genes overlap with the networks of rASD genes. XP-all: union of XPbroad and XP-specific networks; rASD net 1: network from (Chang et al., 2015); rASD net 2: networks 3-5 from (Willsey et al., 2013); rASD net 3: networks 4-6 from (Willsey et al., 2013). **C)** Recurrent DE genes are enriched for brain-specific genes. **D)** Recurrent DE genes are predominantly upregulated across four different cohorts of hiPSC-derived neuronal models of ASD (Methods). **E)** Recurrent DE genes overlap strongly with only the neurodevelopmental network of brain-specific rASD genes. **F)** Effects of damaging mutations in rASD genes converge on the recurrent DE and XP-specific genes. The colors illustrate the odds ratio, and the red borders demonstrate significant odds ratios (FDR <0.1). Data sources: PTEN/MeCP2/FMR1 k/d: Lanz et al., 2013; BAF170 k/d: Tuoc et al., 2013; SOX5 o/e: Parikshak et al., 2016; FMR1 ChIP-Seq: Darnell et al., 2011; CHD8 ChIPSeq 1: Cotney et al., 2015; CHD8 ChIP-Seq 2: Sugathan et al., 2014; CHD8 ChIP-Seq 3: Gompers et al., 2017. k/d: knockdown; k/o: knockout; o/e: overexpression. **G)** Recurrent DE genes are enriched for the core neurodevelopmental processes involved in ASD (nodes in yellow). Brain-specific rASD genes overlap with the recurrent DE genes. Broadly-expressed and some of brainspecific rASD genes regulate the ASD core processes. Effects of many of broadly-expressed rASD genes on the core processes are channeled through the signaling pathways.

Given that the XP-broad and XP-specific genes inter-connect through the signaling pathways, we next examined whether they are enriched for regulators of PI3K/AKT, RAS/ERK, and WNT/*β*-catenin signaling pathways. To do this, we analyzed genome-wide data wherein gene knockouts are screened for their effect on phosphorylation of the signaling pathways in stem cells (Brockmann et al., 2017). We found that XP-broad genes are enriched for the regulators of these three signaling pathways, suggesting an upstream regulatory role for XP-broad genes (Fig 2D-E). However, XP-specific genes are not enriched for the regulators of the signaling pathways (Fig 2D-E).

### Broadly-expressed and brain-specific rASD genes connect differently with ASD molecular perturbations

With the hierarchy of broadly-expressed and brain-specific rASD established, we aimed to unravel the roles of the rASD genes in ASD neurobiology by reanalyzing four transcriptomic studies on hiPSC-derived neurons from individuals with ASD. Three studies were conducted on subjects with idiopathic ASD (Liu et al., 2017; Mariani et al., 2015; Schafer et al., 2019) and one transcriptomic study on hiPSC-derived neurons from a subject with ASD and a SHANK2 loss-of-function mutation (Zaslavsky et al., 2019).

A meta-analysis of the four iPSC studies identified 599 “recurrently” perturbed DE genes in neural progenitor and neuron models of ASD (Fig S6). These recurrent DE genes were mostly brain-specific (OR: 4.56; *P*: 4.35×10^−60^; Fig 3C; Fig S6) and were significantly upregulated in ASD neuron models (Fig 3D). The recurrent DE genes are anticipated to include the core molecular processes underlying ASD. Supporting this, we found a significant overlap between the recurrent DE genes and networks of rASD genes (Chang et al., 2015; Willsey et al., 2013) (Fig 3B). Thus, ASD associated perturbations in neuronal cells converge on genes and processes that are specific to developing neurons.

We next examined the overlap of recurrent DE genes with XP-specific and XP-broad genes, separately. Recurrent DE genes significantly overlap with the XP-specific genes even after controlling for their enrichment for brain-specific genes in general (Fig 3E; OR: 2.29; *P*: 2.0×10^−4^). In contrast, the recurrent DE genes did not overlap with XP-broad genes (OR: 0.30; *P*: 0.027; Fig 3E).

### Intermediate signaling pathways are connected to recurrent DE genes in ASD neuron models

Since XP-broad genes regulate the RAS/ERK, PI3K/AKT, and WNT/*β*-catenin signaling pathways, we tested if the signaling pathways impact recurrent DE gene expression, including the XP-specific network. First, we found the knockdown of three example rASD genes (i.e., PTEN, MeCP2, and FMR1) that regulate the RAS/ERK, PI3K/AKT, and WNT/*β*-catenin signaling pathways significantly impact the recurrent DE genes (Lanz et al., 2013) (Fig 3F). Moreover, consistent with the overlap of recurrent DE genes and the XP-specific genes, the XP-specific genes are also significantly perturbed in response to the knockdown of PTEN and FMR1, but not MeCP2 (Fig 3F and S8). Thus, while the XP-specific network overlaps with the recurrent DE genes, the XP-broad network includes regulators of recurrent DE and XP-specific gene expression.

### Genetic mutations in broadly-expressed and brain-specific rASD genes converge on shared neurodevelopmental processes

Our results suggest that broadly-expressed rASD genes could regulate the recurrent DE genes and brain-specific rASD genes. Moreover, the single cell analysis suggests that the expression of broadly-expressed rASD genes are mostly affected by cell migration from VZ to SVZ and CP. How do specific genetic aberrations in broadly-expressed rASD genes affect recurrent DE and XP-specific genes? We examined the genes and processes perturbed by dysregulation of broadly-expressed BAF170 (Alfert et al., 2019; Tuoc et al., 2013) and brain-specific SOX5 (Parikshak et al., 2016) rASD genes. Although in different gene groups, both genes modulate proliferation and maturation of neurons during migration across the laminae and are functionally associated with the PI3K/AKT, RAS/ERK, and WNT/*β*-catenin signaling pathways (Pfister and Ashworth, 2017)(Kurtsdotter et al., 2017). In the neuronal cells, the BAF170 knockout and SOX5 over-expression yields DE genes that significantly overlap with the recurrent DE genes and XP-specific genes (Fig 3F).

We next analyzed ChIP-Seq data from CHD8 (Cotney et al., 2015; Gompers et al., 2017; Sugathan et al., 2014) and FMR1 (Darnell et al., 2011), two rASD genes that modulate expression of many rASD genes. Interestingly, this analysis suggested that CHD8 and FMR1 differ in the gene groups that they regulate through direct binding (Fig 3F). Specifically, CHD8 preferentially regulates XP-broad genes (OR >2.3; *P* <1.2×10^−8^; Fig 3F), while the targets of the FMR1 gene were significantly enriched for XP-broad (OR: 4.46; *P*: 1.1×10^−15^), XP-specific (OR: 3.8; *P*: 6.3×10^−14^), and recurrent DE genes (OR: 3.8; *P*: 7.1×10^−26^) (Fig 3F). The targets of FMR1 were also enriched for the negative control brain-specific ASDsib genes, while showing a stronger enrichment trend for the XP-specific genes (OR: 1.6; *P*: 0.088).

Collectively, in neurons, the effects of most broadly-expressed and brain-specific rASD genes converge on common core processes in ASD that are neuron-specific. Moreover, the brain-specific rASD genes show strong enrichment with the core ASD processes (Fig 3G).

### Non-neuronal cells harbor dysregulated transcription in broadly-expressed rASD genes

Both neuronal and glial cells are implicated in ASD pathobiology (Ballas et al., 2009; Derecki et al., 2012; Gupta et al., 2014; Kasah et al., 2018; Morgan et al., 2012; Morgan et al., 2010; Suzuki et al., 2013; Voineagu et al., 2011). The XP-broad network is expressed across cell types, while the XP-specific network is enriched in neuronal cells. We hypothesized perturbations of the XP-broad network could dysregulate diverse cell types, while XP-specific network perturbations would mostly impact neurons. Thus we analyzed rASD gene group expression in microglia across developmental stages (Galatro et al., 2017; Matcovitch-Natan et al., 2016; Zhang et al., 2014). XP-broad genes were strongly expressed in microglia, but XP-specific gene expression was mostly limited to neuronal cells (Fig 4A and Fig S9). To further assess the activity of the broadly-expressed network in developing non-neuronal cells, we analyzed its expression during hematopoietic differentiation (Novershtern et al., 2011). We found broadly-expressed rASD genes are strongly expressed (Fig 4B) and their neurodevelopmental network is active during the differentiation of this non-neuronal tissue (Fig 4C). Although brain-specific rASD genes showed low expression during hematopoiesis (Fig 4B), their expression, on average, anti-correlated with broadly-expressed genes (Fig 4D), resembling their neurodevelopmental gene expression correlation patterns.

**Fig 4.**
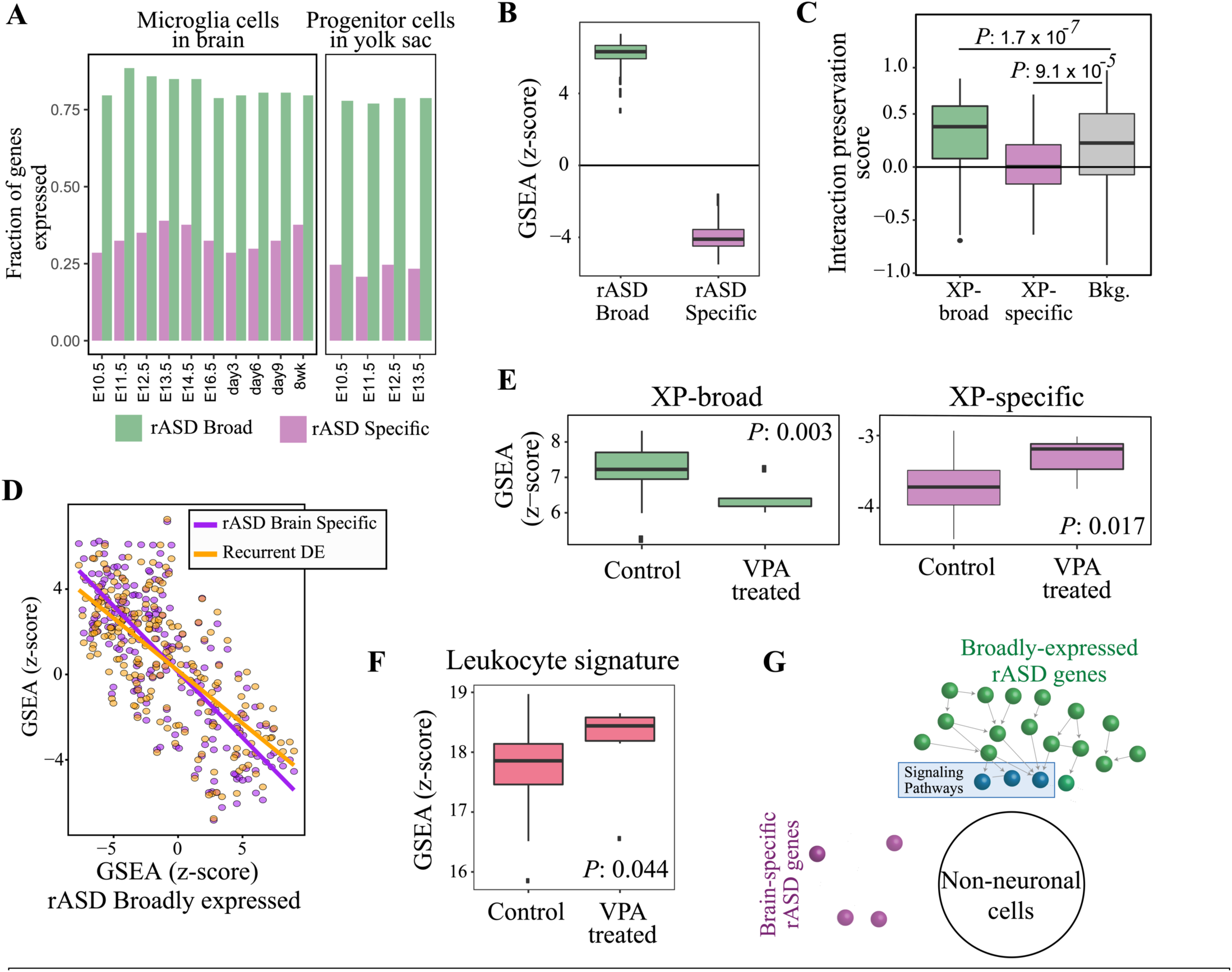
Genetic perturbations in broadly-expressed genes impact non-neuronal cells. **A)** The fraction of broadly-expressed and brain-specific rASD genes expressed in stages of developing mouse yolk sac progenitors and microglia. **B)** Broadly-expressed rASD genes show strong expression during hematopoiesis. The z-scores were calculated by GSEA. **C)** The broadly-expressed gene neurodevelopmental network is active during hematopoietic differentiation. Score magnitude quantifies co-expression strength, and sign signifies if the co-expression direction was preserved (positive indicates co-expression in the same direction during brain development and hematopoiesis; see methods). To avoid bias from preferential expression of the brain-specific genes in the brain, the analysis was limited to the genes that show expression during hematopoiesis differentiation. **D)** Expression of the broadly-expressed rASD genes is anti-correlated with brain-specific rASD and recurrent DE genes during hematopoiesis. Expression of each gene was normalized to have mean of zero and standard deviation of one. GSEA compared rASD gene group expression with all genes expressed during hematopoiesis (Mann-Whitney-Wilcoxon test). A positive/negative z-score indicates that the gene set tends to be up/down regulated compared to background. **E)** The neurodevelopmental gene networks of broadly-expressed and brain-specific genes demonstrate opposite responses to VPA treatment in microglia. Z-scores are comparing GSEA results on the broadlyexpressed and brain-specific rASD genes in microglia with and without VPA treatment. **F)** ASD DE signature in leukocytes is upregulated in VPA-treated microglia. **G)** In non-neuronal cells, the network of broadly-expressed rASD genes are preserved. However, network of brain-specific rASD genes is weakly expressed and is not conserved in non-neuronal cells.

We next examined how non-neuronal cell gene expression relates to the rASD gene network. For this, we leveraged transcriptomics from microglia when treated with Valproic acid (VPA), a modulator of the PI3K/AKT signaling pathway (Cho et al., 2019). Exposure to VPA during pregnancy is associated with increased ASD incidence (Christensen et al., 2013). Corroborating the connection with ASD molecular pathobiology, the DE genes following VPA treatment in microglia are enriched in ChIP-Seq targets of CHD8 and FMR1 (Fig S10). Furthermore, gene set enrichment analysis (GSEA) suggested XP-specific genes are upregulated in response to VPA in microglia (*P*: 0.017; Fig 4E), even though their expression was very low in microglia (Fig 4E). In contrast, the XP-broad genes were down-regulated (*P*: 0.003; Fig 4E), further emphasizing that the two gene groups involve different but potentially connected networks.

We previously identified a DE gene signature in leukocytes that is modulated by rASD genes and more active in ASD individuals (Gazestani et al., 2019). Here we further tested if dysregulation in this gene signature overlaps with the XP-broad or XP-specific genes. While the leukocyte gene signature did not overlap significantly with XP-broad genes nor XP-specific genes (Fig S10), the rASD genes regulating the leukocyte DE signature were predominantly broadly-expressed (OR: 2.31; *P*: 3.5×10^−8^; Fig S10). Furthermore, the leukocyte DE signature significantly overlapped with the DE genes following VPA treatment of microglia (OR: 1.35; *P*: 0.0063; Fig S10). Moreover, the leukocyte signature was upregulated in microglia treated with VPA (*P*: 0.044; Fig 4F), as we observed in ASD subjects with more severe symptoms and in recurrent DE genes in hiPSC-derived neuronal models of ASD (Gazestani et al., 2019).

Together, the network of broadly-expressed rASD genes is conserved across cell types. Therefore, its perturbations could impact non-neuronal cells including microglia and leukocytes (Fig 4G). This may be because the implicated PI3K/AKT, RAS/ERK, and WNT/*β*-catenin signaling pathways are conserved and functional across all cell types and tissues.

### Dysregulation of broadly-expressed rASD genes has more extensive neurodevelopmental effects

Since the XP-broad network is expressed earlier in neurodevelopment, also strongly expressed in non-neuronal cells, and associated with transcriptomic dysregulation in leukocytes of ASD subjects, we hypothesized that the XP-broad network could exert a wider physiological impact by affecting both neuronal and non-neuronal cells.

ASD risk genes overlap substantially with other neurodevelopmental disorders, including developmental delay (DD) and intellectual disability (ID). Interestingly, we found high confidence DD genes are enriched for broadly-expressed genes (OR: 1.36; *P*: 5.7×10^−4^; Fig 5A). Similarly, rASD genes that overlap with DD genes are enriched for broadly-expressed genes (OR: 1.6; *P*: 0.034; Fig 5A).

**Fig 5.**
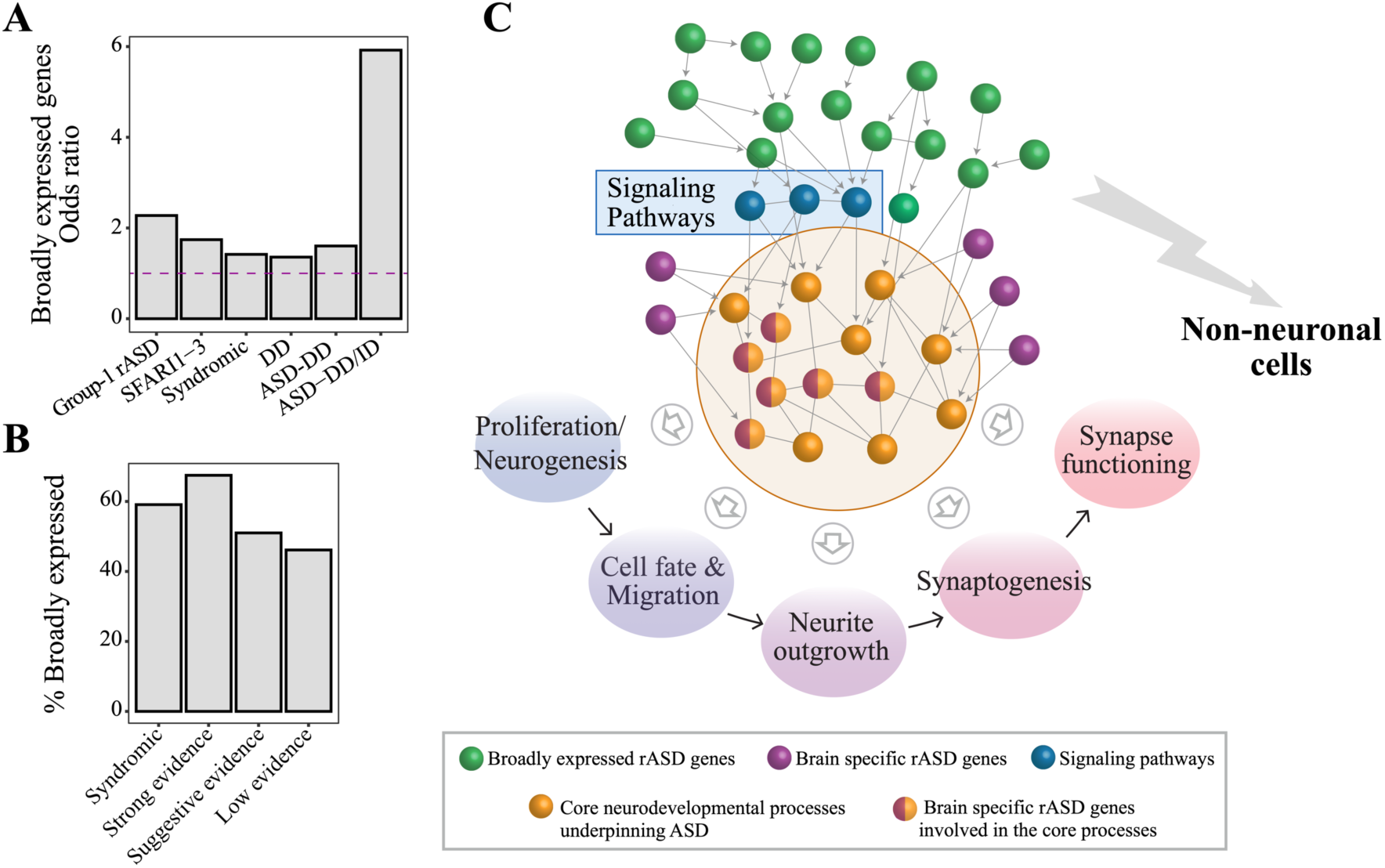
A unifying genetic-transcriptomic model of ASD. **A)** Broadly-expressed ASD risk genes exert large impact on brain development. Genes implicated in DD/ID, syndromic, and high confidence rASD genes are enriched among broadly-expressed genes compared with lower confidence rASD genes. The Y-axis quantifies relative enrichment of each gene group for broadly-expressed genes compared with all currently implicated rASD genes except for the DD group. The enrichment of DD genes was based on the analysis of all genes expressed during prenatal and early postnatal brain development. ID: intellectual disability; DD: developmental delay; Group 1: rASD genes whose mutations are not identified in a large cohort of healthy individuals based on the ExAct database (Kosmicki et al, 2017); ASD-DD: rASD genes overlapping with the DD risk genes from the DDG2P database; ASD-DD/ID: Genes overlapping between Group 1 rASD genes and DD/ID risk genes from (Kosmicki et al, 2017). **B)** Fraction of broadly-expressed genes in different rASD gene categories from the SFARI database. **C)** Schematic illustration showing the model of the genetic architecture of ASD articulated in this study. rASD genes are composed of the two groups of broadly-expressed and brainspecific genes (Fig 1B). Broadly-expressed rASD genes are involved in gene expression regulation, cell proliferation, and cell fate determination (Fig 3A). In neuronal cells, the effect of broadly-expressed rASD genes converge on the ASD neurodevelopmental core processes spanning proliferation and neurogenesis to synaptogenesis and synapse functioning (Fig 3A & 3F). Importantly, the impact of many broadly-expressed rASD genes is channeled through the RAS/ERK, PI3K/AKT, and WNT/*β*-catenin signaling pathways (Fig 2D). Brain-specific rASD genes overlap with the ASD core processes, while some regulate these core processes as well (Figs 3C-F). Here, we speculate that the recurrent DE genes from the four independent iPSC studies on ASD are highly enriched for the core processes underpinning the disorder (represented by the yellow nodes). Finally, perturbations to the broadly expressed genes are poised to impact non-neuronal cell and could underliediverse comorbidities in ASD (Figs 4E-F & 5A).

ASD shows co-morbidity with other disorders and many rASD genes are categorized as “syndromic”, since their mutations are associated with additional characteristics and dysmorphologies that are not required for an ASD diagnosis (Abrahams et al., 2013). Syndromic rASD genes are enriched for the broadly-expressed genes (OR: 1.42; *P*: 0.031; Fig 5A), thus further supporting the possibility that broadly-expressed genes may have broad impact on neurodevelopment.

By comparing genetic information from ASD and a large cohort of healthy individuals, an independent study found that many rASD genes are also mutated in healthy individuals and only a relatively small subset occurs exclusively in ASD (Kosmicki et al., 2017). Consistent with the hypothesis that broadly-expressed rASD genes would have earlier and more severe phenotypes, we found broadly-expressed genes are enriched among rASD genes that are not mutated in healthy individuals (OR: 2.28; *P*: 9.23 × 10^−7^; Fig 5A). The enrichment further increases for broadly-expressed rASD genes that overlap with DD and ID genes (OR: 5.9; *P*: 2.28×10^−4^; Fig 5A). Likewise, rASD genes with high confidence SFARI scores of 1 to 3 were more broadly-expressed compared to rASD genes with SFARI scores of 4 to 6 (OR: 1.74; *P*: 0.007; Fig 5). Collectively, these results demonstrate that although rASD genes are enriched for the brain-specific genes, most recognized high confidence rASD genes are broadly-expressed and their mutations could potentially have a larger impact on neurodevelopment (Fig 5C).

## DISCUSSION

ASD has a complex and heterogenous genetic makeup where perturbations to many brain-specific and broadly-expressed genes underlie its heritability. The question has remained on how these diverse genes are organized to result in common ASD traits. Genetic theories, such as the oligogenic, polygenic, and omnigenic models, provide invaluable insights into how the genetics of complex diseases could be explained by tissue-specific and broadly-expressed genes. Importantly, these models propose that genetic aberrations for both tissue-specific and broadly-expressed risk genes could converge through gene regulatory networks onto neurodevelopmental processes (Boyle et al., 2017; Califano and Alvarez, 2017; Courchesne et al., 2020; Gazestani et al., 2019). Here we systematically tested this concept for rASD genes. We found the heterogenous rASD genes form two major groups of broadly-expressed and brain-specific genes. In contrast to the prevailing view, which assumes the rASD genes form a coherent network as a single group, we found that broadly-expressed and brain-specific genes form separate networks that are connected through gene regulatory networks, such as the PI3K/AKT, RAS/ERK, and WNT/*β*-catenin signaling pathways. The two networks function successively during prenatal neural development, with broadly-expressed rASD genes involved in cell proliferation, transcriptional regulation, and cell fate commitment, while brain-specific rASD genes contribute more to neural maturation, and synapse formation and function. Thus, broadly-expressed rASD genes perturb brain-specific rASD genes and recurrent DE genes in ASD through gene regulatory networks (Fig 5C). Although the involvement of rASD genes in different gene networks has been proposed (Courchesne et al., 2020; Parikshak et al., 2013; Pinto et al., 2014; Satterstrom et al., 2020; Yuen et al., 2017; Zhou et al., 2019), our study describes how they form a hierarchical network converging on neurodevelopmental roles providing a clear view on the genetic architecture of ASD and insights into its progression.

The biological differences of broadly-expressed and brain-specific rASD genes are rooted in different transcriptional programs. Broadly-expressed genes show peak expression in neural progenitor cells and their expression gradually decreases during transitions from VZ to SVZ and CP in both excitatory and inhibitory neurons at early-to-mid prenatal ages. However, brain-specific rASD gene expression shows differential regulation at cell type transitions with minimal expression in neural progenitor cells and peak expression in mature excitatory and inhibitory neurons. Brain-specific rASD genes also show a laminae-dependent expression pattern, with lowest expression in VZ. The temporal expression of the two rASD groups highlights the importance of neurodevelopmental processes during the second and third trimesters in ASD. At this time the expression of the two gene categories transition, thus marking a critical time for ASD. At this phase, neural processes transition from proliferation to neurogenesis, cell fate determination, migration, and maturation (Courchesne et al., 2019; Li et al., 2018; Sahin and Sur, 2015; Willsey et al., 2013). Dysregulation of these interconnected processes can result in unbalanced generation of neuronal types that are premature with aberrant growth and synaptogenesis (Courchesne et al., 2019; Fang et al., 2014). Furthermore, these perturbations could upset the timing of neurodevelopmental stages responsible for the correct organization of neural circuits at important initial stages. For example, the surge in synaptogenesis at the late prenatal and early postnatal ages needs to be followed by extensive, experience-based synapse pruning processes. However, an early up-regulation of the brain-specific genes, as observed in hiPSC-derived neuron models of ASD, could trigger the formation of synapses at a time that brain does not receive environmental inputs, further compounding the aberrant formation of neural circuit organization.

Core molecular processes underlying ASD are expected to be commonly perturbed in many subjects with ASD (Courchesne et al., 2020; Liu et al., 2019). Our recurrent DE gene analysis found shared transcriptomic perturbation patterns across four independent ASD cohorts with diverse genetic backgrounds. These core perturbed processes include neurogenesis, neural migration, maturation, and synapse functioning. These shared gene expression patterns overlapped with the network of brain-specific rASD genes (i.e., XP-specific genes) and are predominantly upregulated in ASD neurons (Fig 5C). However, the perturbed broadly-expressed rASD genes impact both XP-specific and recurrent DE genes, showing broadly-expressed rASD genes are key regulators of ASD transcriptional programs. Since recurrent DE genes were also perturbed in idiopathic ASD subjects, our results support existence of common molecular perturbations between ASD common variants and rASD genes. Furthermore, the lack of overlap between broadly-expressed and recurrent DE genes suggest that ASD individuals can have perturbations in a wider range of broadly-expressed genes. Therefore, on average, broadly-expressed genes do not show strong perturbations, but specific changes in a broadly-expressed gene can underlie ASD in individuals. This model suggests that although broadly-expressed genes contribute to ASD liability, the dysregulation of brain-specific genes including XP-specific and recurrent DE genes is necessary and ultimately impact neural development in ASD (Fig 5C).

Our results point to at least two genetic subtypes of ASD. While the broadly-expressed and brain-specific rASD genes converge on shared neurodevelopmental processes, both show group-specific characteristics with possibly differing influences on brain development. The broadly-expressed genes express earlier during brain development, while the brain-specific gene network remains active throughout life. Moreover, broadly-expressed genes are active in glial (e.g., microglia) and other non-neuronal cells (e.g., leukocytes). Thus, broadly-expressed genes could directly impact non-neuronal cells throughout life, but their impact on neurons seems mostly limited to early neurodevelopment. The full clinical implications of these findings remain to be established. However, our results suggest broadly-expressed rASD genes exert larger impacts on brain development. Corroborating these results, another study recently identified an enrichment of ID risk genes in radial glial cells compared to rASD genes (Polioudakis et al., 2019). The broadly-expressed risk genes might also explain the common molecular perturbations observed between ASD and its non-brain related comorbidities (Nazeen et al., 2016). Further analysis of clinical and treatment data in the context of our genetic architecture could elucidate possible subtype specific traits. This paradigm could also clarify functional genomic analyses by partitioning subjects into molecular subtypes of ASD and enable the wider use of peripheral cell types that are easier to obtain from children (e.g., blood and skin fibroblasts).

Finally, the seemingly enormous heterogeneity of ASD-relevant genes has stymied a concise understanding of how they affect ASD neural development (Courchesne et al., 2019; de la Torre-Ubieta et al., 2016; Pinto et al., 2014). The unifying model presented here builds on the previous genetic models that emphasize small additive effects of numerous genes (Turner et al., 2017; Weiner et al., 2017), but goes beyond by explaining how the many specific rASD genes cause prenatal maldevelopment. Our model goes beyond single gene models by placing each gene in the context of all other rASD genes. Thus, this genetic model provides a clearer picture into how and when each risk gene contributes to neurodevelopment in ASD. In sum, classifying rASD genes into their tissue specificity provides a much clearer picture on the organization of the genetic makeup of ASD and a genomic foundation to analyze and interpret the temporal and developmental role of each gene. This model opens avenues for more precise etiological and treatment research on the cell biology of living ASD infants, children and adults.

## ACKNOWLEDGEMENTS

The authors also thank Dr. Evan Boyle for input in this work. This work was supported by National Institute of Mental Health (NIMH) R01-MH110558 (E.C., N.E.L.), National Institute on Deafness and Other Communication Disorders (NIDCD) R01-DC016385 (E.C.), and generous funding from the Novo Nordisk Foundation through the Center for Biosustainability at the Technical University of Denmark (NNF10CC1016517 N.E.L.).

## METHODS

### Broadly-expressed and brain-specific genes

Genes expressed in the brain cortex were categorized into the two groups of broadly-expressed and brain-specific based on their gene expression patterns across human tissues. Specifically, we downloaded transcript per million (TPM) normalized gene expression from 10,259 samples across 26 tissues from GTEx portal (Consortium, 2015). In addition to brain and nerve, the dataset included transcriptome data from 24 non-neuronal tissues, including: Adipose, Adrenal Gland, Blood Vessel, Breast, Blood, Skin, Colon, Esophagus, Heart, Liver, Lung, Salivary Gland, Muscle, Ovary, Pancreas, Pituitary, Prostate, Small Intestine, Spleen, Stomach, Testis, Thyroid, Uterus, and Vagina. We next defined a gene expressed in a tissue if met two criteria. First, the gene TPM expression level was ≥ 3 in at least half of the samples from the tissue. Second, the median expression of the gene was equal or larger than its 25-percentile expression in GTEx cortex samples. The second criterion was included to account for the differences in the base expression level of the genes and their dosage dependent translation and function. Broadly-expressed genes were defined as genes that were expressed in ≥ 50% of non-neuronal tissues (i.e., tissues other than brain and nerve).

The broadly-expressed and brain-specific genes included genes that were expressed in the adult cortex based on GTEx dataset. The primary rational for this procedure was that if an rASD gene is not expressed in the adult cortex tissue, its expression pattern in other adult tissues may not be a reliable indicator of its function either. This procedure resulted in the exclusion of 17% of the 232 rASD genes from our main analysis. A possible bias of this procedure towards the selection of neuron specific genes is mitigated by the fact that brain tissue is composed of both neuronal and non-neuronal cell types. Nevertheless, we investigated if the excluded genes show evidence of bias in either broadly-expressed or brain-specific genes. For this, we leveraged our findings from on the neurodevelopmental gene expression pattern differences of the two gene groups. Specifically, the examination of the neurodevelopmental gene expression patterns indicated that the excluded genes are randomly distributed between the genes with early and late peak expression patterns during the prenatal and early postnatal neurodevelopmental stages (Fig S2). We also found that excluded genes are significantly connected to the neurodevelopmental networks of both broadly-expressed and brain-specific rASD genes. Since the neurodevelopmental networks of broadly-expressed and brain-specific genes are separated, the association of the excluded genes with both networks further supports that these excluded genes are a mix of the two gene groups (Fig S2). Although it was possible to assign categories to the genes with no expression in the adult cortex based on the available functional annotation data and their gene expression patterns during the prenatal and early postnatal brain development, we opted to exclude these genes to facilitate more systematic analysis and avoid biases due to the judgements. Moreover, the employed systematic procedure empowered us to include proper and well controlled background genes in our downstream analyses.

### Mouse tissue gene expression data

Processed gene expression data were downloaded from ENCODE project web portal (Pervouchine et al., 2015). Gene expression data were merged across the biological replicates. In each sample, genes with non-parametric irreproducibility ascertainment (npIDR) <0.1 and merged RPKM >1 were marked as expressed (Pervouchine et al., 2015). To evaluate the correspondence of broadly-expressed and brain-specific assignments of rASD genes from GTEx human tissue data in mouse, we next examined the expression of rASD genes in 20 non-neuronal mouse tissues including: Adrenal gland, Duodenum, Stomach, Small Intestine, Ovary, Testis, Heart, Kidney, Lung, Thymus, Mammary Gland, Spleen, Colon, Liver, Genital Adipose, Subcutaneous Adipose, Large Intestine, Placenta, Limb, and Urinary Bladder.

### ASD (rASD), intellectual disability (ID), and developmental disorder (DD) risk genes

We focused on rASD genes with a higher probability of relevance to ASD etiology. To confirm the robustness of the results, we also considered two different, yet overlapping gene sets, defined based on different criteria. The primary gene set was composed of 232 rASD genes that were supported by at least two different resources based on information available in SFARI database in June 2019 (Abrahams et al., 2013). The secondary gene set was composed of high and strong confidence SFARI genes (SFARI score 1 and 2 as of January 2020 version of the database) and 102 risk genes recently identified as significant (FDR <0.1) in the largest exome-sequencing study of ASD to date (Satterstrom et al., 2020). The secondary gene set of rASD genes was composed of 385 unique risk genes. Unless otherwise noted, we reported results that were reproducible in both rASD gene sets. The main figures are based on the primary gene set and the confirmation of the results from the secondary set is presented in the supplementary material. Previously published genes with likely gene damaging and synonymous mutations in siblings of subjects with ASD, who developed normally (ASDsib genes) were retrieved from (Iossifov et al., 2014). ASDsib genes were used as negative controls to ensure the specificity of the results reported hereafter to rASD genes (Supplementary Figures). Gene names in these datasets were converted to Entrez IDs using a combination of DAVID and BioMart tools.

We also tested if broadly-expressed and brain-specific rASD genes demonstrate preferential enrichment among syndromic, Group 1 (Kosmicki et al., 2017), SFARI categories 1-3, and overlapping genes between ASD and DD/ID genes. Syndromic rASD genes were downloaded from SFARI database on March 15, 2020. Group 1 rASD genes were retrieved from (Kosmicki et al., 2017) and defined as the risk genes with pLI score of above 0.9 and de novo protein truncating variants in individuals with ASD. High confidence DD risk genes were downloaded from the DDG2P database on Aug. 15, 2019 (Bragin et al., 2014). A secondary list of high-confidence DD/ID genes that overlap with ASD were downloaded from the list of group 1 genes from (Kosmicki et al., 2017).

### Neurodevelopmental gene expression patterns of ASD risk genes

Normalized RNA-Seq gene expression patterns of the rASD genes were retrieved from the BrainSpan portal (BrainSpan, 2016; Kang et al., 2011). To isolate effects that are characteristics of the broadly-expressed and brain-specific gene groups in general from those of rASD gene groups, we first categorized all genes expressed in the developing brain as brain-specific or broadly-expressed based on their tissue gene expression patterns using a similar approach to that employed for rASD genes. We next tested if, in general, broadly-expressed and brain-specific genes show differences in their neurodevelopmental gene expression patterns based on the neocortex RNA-Seq data from the BrainSpan. To do so, for each gene, we calculated the correlation of its gene expression pattern with the neurodevelopmental time point (modeled as an ordinal variable) between 8 pcw to 2 years after birth using the robust biweight midcorrelation efficiently metric implemented in WGCNA (Langfelder and Horvath, 2012). Interestingly, this analysis revealed that the majority percentage of broadly-expressed genes have peak expression during early prenatal development and then downregulated during later prenatal and early postnatal brain development as indicated with the negative correlation of their expression patterns with the neurodevelopmental time points (Fig 1F). On the contrary, a majority of brain-specific genes show a gradual upregulation pattern during prenatal development with peak expression at the late prenatal and early postnatal time periods (Fig 1F). We next examined the gene expression pattern of broadly-expressed and the brain-specific rASD genes during the neurodevelopment. Interestingly, this analysis indicated that the rASD genes show significantly different gene expression patterns compared to the overall broadly-expressed and the brain-specific genes (Fig 1E-F).

Brain-specific rASD genes exhibited a bimodal gene expression pattern (Fig 1F). Therefore, We divided the brain-specific rASD genes to two groups of “early” and “late” brain-specific rASD genes based on the correlation of their expression patterns with the developmental time points in neocortex from (8pcw to 2yrs) using the robust biweight midcorrelation efficiently metric implemented in the WGCNA package (Langfelder and Horvath, 2012) and treating the time point as an ordinal variable. Genes whose expression positively correlated with the developmental time points were termed as late brain-specific genes. Similarly, genes whose expression negatively correlated with the neurodevelopmental time points were termed as early brain-specific genes.

### Neurodevelopmental networks of ASD risk genes

To examine the connection of rASD genes with one another and with molecular processes related to brain development, we constructed a neurodevelopmental gene network composed of all genes expressed during brain development using the normalized RNA-Seq gene expression data from the BrainSpan resource (BrainSpan, 2016; Kang et al., 2011). Briefly, we first retrieved strong evidence physical and regulatory interactions from the Pathway Commons database (Cerami et al., 2011) and updated the network to include interactions from the most recent Reactome (Fabregat et al., 2016) and BioGrid (Chatr-Aryamontri et al., 2017) databases. To make the network relevant to neurodevelopment, we pruned the network to retain only interactions between strongly co-expressed gene pairs in the neocortex. A gene pair was deemed as strongly co-expressed if their co-expression strength was ≥ 0.5 as judged by unsigned Pearson’s correlation coefficient metric. Considering the prenatal and early postnatal onset of ASD, RNA-Seq brain gene expression data from 8pcw to 1yr was considered to assess the co-expression of interactions. To calculate correlations, normalized RPKM gene expression values were log2(x+1) transformed. Furthermore, since the interactions of highly connected genes are not informative, we excluded the top 5% highly interacting genes from the neurodevelopmental gene network. We next extracted the neurodevelopmental sub-networks of broadly-expressed and brain-specific rASD genes by considering interactions that involved these rASD genes.

### Examining the within group interaction density of broadly-expressed and brain-specific rASD genes

To test if broadly-expressed and brain specific rASD genes are significantly connected to common gene neighbors and hence form a dense network, we compared the node degree distribution of the neighbors in the broadly-expressed and brain-specific rASD subnetworks with background distributions. To construct the background distributions for the broadly-expressed rASD network, iterating 1000 times, we randomly selected a gene set of broadly-expressed non-rASD genes with the same node degree distribution as the broadly-expressed rASD genes ±20% and examined the node degree distribution of their neighbors. For each background distribution, a *P* was estimated by examining the node degree distribution of neighbors between the observed (i.e., the neighbors of broadly-expressed rASD genes) and expected by chance (i.e., random gene sets of broadly-expressed non-rASD genes) using the Wilcoxon-Mann-Whitney test. The *P* was next z-score transformed using the coin package in R. The collection of 1000 z-scores (represented as a boxplot in Fig 2A) were next tested to identify if they show a significantly skewed distribution or are randomly distributed around zero. A similar procedure was used to examine the within group interactions of brain-specific rASD genes.

### Examining the between group interaction density of broadly-expressed and brain-specific rASD genes

To test the density of between-group interactions of broadly-expressed and brain-specific rASD genes, iterating 1000 times, we randomly selected a gene set of broadly-expressed and brain-specific non-rASD genes with the same node degree distribution as their rASD counterparts ±20%. We next examined the number of common interactions between the two random gene groups. An empirical *P* was estimated by comparing the observed number of common interactions between the two rASD gene groups with the background distribution from the randomized gene sets (Fig 2B).

To ensure that the network level separation of broadly-expressed and the brain-specific rASD genes is not due to a bias in the current state of knowledge of the human interactome, we constructed two additional full co-expression networks of neurodevelopmental stages by considering only gene pairs with the correlation strength (i.e., unsigned Pearson’s correlation coefficient) of 0.5 and 0.7 based on the BrainSpan RNA-Seq data from neocortex (BrainSpan, 2016; Kang et al., 2011), separately. We evaluated the between-group interaction patterns of rASD genes in these two co-expression networks using the same empirical approach elaborated above and found a significant separation of the broadly-expressed and brain-specific genes in these co-expression networks as well (Fig S5).

A strong co-expression of rASD genes at 12 to 24 pcw has been reported previously (Willsey et al., 2013). Therefore, we tested if the brain-specific and broadly-expressed rASD genes become strongly interconnected during these neurodevelopmental stages. We constructed a co-expression network from the neocortex RNA-Seq data during 12 to 24 pcw (unsigned correlation >0.7). We next tested the extent of common gene neighbors between the broadly-expressed and the brain-specific genes using the method elaborated above. Similar to the other constructed gene networks, we found the broadly-expressed and brain-specific genes are significantly depleted for common gene neighbors at these neurodevelopmental stages as well (Fig S5).

### Identifying significant network neighbors of rASD genes

We employed network propagation concepts to identify genes that are significantly connected to rASD genes (Vanunu et al., 2010). In these semi-supervised approaches, information on the annotated genes (i.e., gene labels indicating whether a gene is an rASD or ASDsib) are diffused through gene networks to identify the relevant genes that are significantly connected to them. The significance of a neighbor depends on the number of interactions of both the labeled genes and the neighboring gene as encoded in the network structure. Our main analysis of rASD genes included genes with a higher confidence of relevance to ASD, constituting only a subset of currently implicated risk genes. Network propagation techniques are especially well suited for the purpose of our analyses as they allow to incorporate into the analysis the available knowledge on hundreds of lower confidence risk genes that show de novo damaging mutations in individuals with ASD.

We performed network propagation on the neurodevelopmental gene network from prenatal (starting 8 pcw) to early postnatal (ending at 1 years of age) stages. The neurodevelopmental network was represented with an adjacency matrix *W*. In the *W*, *w*_*ij*_ is one if gene interacts with gene *j*, and *w*_*ij*_ is zero otherwise. We performed a network propagation approach that simulates a random walk with restarts. Specifically, we updated the label of each gene in the network by iteratively combining its label with its network neighbors:

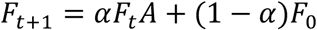

Where *F*_*i*_ represents the gene labels at the time *i*. *F*_0_ represents the initial label of the genes. *A* represents the degree-normalized adjacency matrix of the neurodevelopmental gene network. The matrix *A* was generated by degree-normalizing the adjacency matrix, *W*, through dividing each *w*_*ij*_ by the square root of the product of the node degrees of genes *i* and *j*. In the above formula, *α* is a hyperparameter and specifies the amount the information that genes retain at each iteration. We performed the propagation until the program converged; i.e., the maximum change in the gene labels was less than 10^−5^.

To select the optimum value of *α* hyperparameter, we first assigned the 20% of labeled genes as the held-out set. We next performed a five-fold cross validation using the remaining 80% of gene labels. Specifically, iterating 5 times, we randomly set the label of 20% of rASD and ASDsib genes as unknown (i.e., the score of −0.01; see below). We next performed the propagation and evaluated the accuracy of labels of the 20% genes. We tested the performance of the label propagation using the three *α* values of 0.5, 0.6, and 0.7. The cross-validation results suggested 0.7 as the optimum value of *α*. However, the results were mostly consistent with the α value of 0.6. After the cross validation, the high performing model was tested on the held-out genes.

We performed the network propagation on the two high-confidence rASD gene sets, separately. The label of each gene was represented with a score ranging [-1,1] with score 1 indicating the gene is a high confidence rASD gene and score −1 indicating the gene is an ASDsib gene. We also incorporated the additional available knowledge on ASD genetics in the propagation approach by setting the labels of rASD genes not included in the high confidence set as 0.25. We also assigned a slight penalty to genes other than rASD by setting their labels as −0.01. In our cross-validation experiments, we found that the assignment of slight negative scores to genes with unknown labels improves the propagation results compared to the assignment of zero as the labels.

In our approach, after the convergence of the propagation algorithm, a gene score of above zero suggest that the gene is densely connected to the rASD genes. To assess the significance of the gene scores, iterating 1000 times, we randomly rewired the network while preserving the node degrees. For each gene, an empirical *P* of the gene score was estimated by counting the number of times that the gene obtained the same or higher propagation scores in the shuffled networks compared to its score in the neurodevelopmental gene network. To select the significantly connected genes with the high confidence rASD genes, we only considered the genes that are directly linked to at least two high confidence rASD genes in the neurodevelopmental gene network. Multiple testing on empirical *P* was corrected using the Benjamini-Hochberg procedure. Genes with an FDR <0.1 and a propagation score >0 were deemed as significantly connected to the rASD genes.

The labels of the held-out genes were set to unknown (i.e., score of −0.01) during the cross validation. To assess the performance of the network propagation model, we compared the propagation results on the 20% held-out genes with their known labels. This analysis indicated a strong and significant enrichment of known rASD genes among the genes with FDR <0.1 and score >0 with OR of 6.02 (*P*: 0.0009) and 6.68 (P: 8.8×10^−05^) for the primary and secondary rASD gene sets, respectively. Importantly, this analysis identified only one ASDsib gene, ITPR1, significantly associated with the primary rASD gene set. ITPR1 is a known modulator of PI3K/AKT signaling pathway and it is considered as an rASD gene with a score of 3 in the Feb. 21, 2020 version of the SFARI database. The propagation did not identify any ASDsib genes as significantly associated with the secondary rASD gene set. Network propagation on ASDsib genes did not identify any gene significantly connected to the ASDsib genes (data not shown).

### Evaluating the preservation of rASD neurodevelopmental gene networks in adult cortex

We assessed the preservation of neurodevelopmental networks of rASD genes in adult brain cortex using available normalized expression data from GTEx consortium where gene expression data were corrected for the genotype of individuals, hidden confounding variables, and gender (Consortium, 2015). We employed a gene-centric approach to examine the preservation of the neurodevelopmental networks to account for variations in the number of interactions among rASD genes in the networks. Specifically, for the rASD genes with nine or more interactions in the neurodevelopmental network, we compared the co-expression strength of interactions of each rASD gene in the adult cortex with a background distribution using the Kolmogorov-Smirnov test. For those rASD genes with less than nine interactions, to increase statistical power, we randomly grouped every three of them and compared their interactions collectively with the background distributions. *P* values were next z-transformed for each rASD gene. Finally, for each of the neurodevelopmental networks of broadly-expressed and brain-specific, we tested if the z-scores of their genes are significantly skewed towards positive (i.e., conserved) or negative (i.e., disrupted) using a secondary one-way permutation test with Monte Carlo approximation using the coin package in R. This gene-centric analysis was designed to ensure that the results are not biased by the number of interactions that each rASD gene has in the neurodevelopmental networks. To construct the background distribution for each rASD gene, we randomly selected 20 genes with 1) the same node degree distribution as the rASD genes ±20% and 2) the same co-expression strength with their gene neighbors at neurodevelopmental time point as the rASD gene as examined by the Kolmogorov-Smirnov test (*P* >0.1).

### Gene set overlap analyses

Overlaps between gene sets were examined using the Fisher’s exact test. rASD genes have peak expression in the brain during the prenatal and early postnatal time points (Courchesne et al., 2019; Parikshak et al., 2013). Therefore, to properly control for the background genes for the statistical tests, we used all protein-coding broadly-expressed and brain-specific genes that are expressed during neurodevelopment (based on neocortex RNA-Seq transcriptome data from BrainSpan) as background genes for broadly-expressed and brain-specific rASD genes, respectively. The background genes for the recurrent DE genes were all protein-coding genes expressed in neocortex during the prenatal and early postnatal neurodevelopmental periods. Several comparisons of empirical *P* calculated by permutation tests indicated high concordance rates with the *P* from Fisher’s exact test. To further ensure that potential biases in the rASD genes are not driving the overlap results, we included the genes identified as mutated in typical siblings of subjects with ASD (ASDsib genes) as negative controls in our analyses (Supplementary Figures).

### Gene ontology enrichment analysis

We examined the enrichment of the rASD neurodevelopmental networks and recurrent DE genes for Gene Ontology biological processes (GO-BP) using Fisher’s exact test. Each gene set was tested independently for GO-BP terms with the 10-2000 annotated genes. The terms with FDR <0.1 were next clustered based on their similarity and gene content overlap according to the Amigo GO-BP tree retrieved by the RamiGO package in R. The general terms with more than 1000 annotated genes that spanned two or more clusters were removed. We compared the enrichment patterns of early and late brain-specific rASD networks by focusing only on GO-BP clusters that are in overall enriched in the brain-specific rASD networks as one group. We next compared the odds ratio of the enrichment of each GO-BP cluster between the early and late brain-specific rASD neurodevelopmental networks.

### Human neural progenitor differentiation data

Microarray transcriptome data from the differentiation of primary human neural progenitor cells to neuronal cells were downloaded from the NCBI GEO database (GSE57595). The data were already quantile normalized and ComBat batch-corrected. For genes with multiple probes, we retained the probe with the highest mean expression value (Stein et al., 2014).

### Comparative analysis of the DE genes reported in the four hiPSC-derived neuronal models of ASD

Four hiPSC studies on ASD subjects were considered for this analysis. The list of DE genes were retrieved from *Mariani et al.* (idiopathic ASD with macrocephaly; Genes with FDR <0.1 in Day 11 or Day 31 of hiPSC differentiation to neurons), *Liu et al.* (idiopathic ASD; FDR <0.05 and Fold change >2), and *Zaslavsky et al.* (ASD and SHANK2 loss of function mutation; Genes with FDR <0.1 and Fold change >1.5) studies. We also processed the raw fastq files from the *Schafer et al*. study and considered genes with FDR <0.1 as DE (see below). Background genes were defined as genes expressed in the neocortex during the prenatal and early postnatal stages. Pair-wise overlap analysis of DE genes between the four datasets were conducted by Fisher’s exact test.

### Meta-analysis of the four hiPSC studies on ASD subjects

To identify the connection of rASD genes with molecular perturbations associated with ASD, we considered four independent hiPSC studies, three from idiopathic cases (Liu et al., 2017; Mariani et al., 2015; Schafer et al., 2019) and one from a subject with ASD and a SHANK2 loss-of-function mutation (Zaslavsky et al., 2019). Initial analysis of the four studies demonstrated significant overlap of the DE genes between each pair of the hiPSC studies, suggesting existence of common molecular perturbations in these independent cohorts (see above). However, these studies are each composed of small sample sizes, and the well-known technical variation associated with the iPSC studies could further lower the signal to noise ratio in each dataset.

We performed a meta-analysis of the four studies to identify the genes that show high transcriptional perturbation across the four studies. First, as explained below, we conducted DE analysis at neural progenitor and neuronal stages in each dataset using a standardized analytic pipeline. Second, we combined the *P* of each gene across the four studies using the Stouffer’s z-score method. The *P* were corrected for multiple testing using the Benjamini-Hochberg procedure. In addition to FDR <0.05, we required the DE genes to show a nominal significance (*P* <0.05) in at least two out of four datasets. This resulted in the identification of 599 DE genes that we call “recurrent DE” genes. Examination of the *P* distribution of the recurrent DE genes in each of the four studies demonstrated they are significantly skewed towards smaller *P* compared to the background in all four studies (Fig S6). The overall gene expression pattern of the genes (i.e., up or down regulation) across the four studies were estimated by first averaging the number of the times that each gene was up or down regulated across time points in one study with the average of the 0.5 indicating that the gene is equally up or down regulated across the time points of a study. We next averaged these scores across the four studies to obtain the overall gene expression pattern of each gene.

To further confirm the robustness of the results, we reproduced the presented overlap analyses using a secondary meta-analysis of the four hiPSC studies by considering the genes that are originally identified as DE in at least two out of the four studies (with the exception of *Schafer et al., 2019* study in which we identified the DE genes as described below). The secondary list of DE genes was next refined by considering only those that are connected to at least one more DE gene in the neurodevelopmental gene network. this secondary analysis resulted in the identification of 398 genes as recurrent DE genes. Results on the primary DE genes hold in this secondary list (data not shown).

#### DE analysis of hiPSC study 1 (Liu et al., 2017)

This dataset includes microarray gene expression data from 3 male individuals with ASD and their male siblings. Microarray data were retrieved from NCBI GEO (GSE65106) and quantile normalized. Genes with a log transformed expression level below 4 in more than 50% of samples were excluded. For genes represented with multiple probes, we retained the probe with highest mean expression level and removed the others. Genes differentially expressed across neural progenitor or neuronal stages were identified using the limma package with moderated F-statistic (Ritchie et al., 2015).

#### DE analysis of hiPSC study 2 (Mariani et al., 2015)

This dataset contains RNA-Seq data from 4 male individuals with ASD and macrocephaly and their fathers as controls. We retrieved the fastq sequencing data from SRA database (SRP047194). Reads were aligned to the Genome Reference Consortium GRCh38 using STAR. We excluded the ALT, HLA, and Decoy contigs from the reference genome sequence prior to the alignment. The read counts were estimated using the HTSeq (Anders et al., 2015) and Gencode annotations V26. The Gencode GTF file was collapsed by: 1) removing exons labeled as ’retained_intron’; 2) merging overlapping exons related to the same gene (Consortium, 2015). Gene expression counts related to replicate clones were added together. Low expressed genes were removed using filterByExpr() function in the limma package (Ritchie et al., 2015). Gene expression counts were normalized using the trend approach in limma package. Genes were next examined for the differential expression at days 11 and 31 with the limma package with moderated F-statistics and individuals’ families as a blocking variable.

#### DE analysis of hiPSC study 3 (Schafer et al., 2019)

This dataset includes RNA-Seq data from different time points during the neural differentiation form hiPSCs of 8 male individuals with ASD and macrocephaly and 5 male individuals with TD. We retrieved the fastq data from EMBL-EBI ArrayExpress (E-MTAB-6018). Gene expression counts were estimated using the same procedure as *hiPSC study 2*. After removing low expressed genes and normalization of the data, Genes were examined for the differential expression across neural progenitor differentiation timepoints (i.e., Day 0, 2, 4, 7, and 14) using the limma package with moderated F-statistics.

#### DE analysis of hiPSC study 4 (Zaslavsky et al., 2019)

This dataset includes RNA-Seq data from hiPSC cell lines of a male ASD subject with SHANK2 loss-of-function mutation and its CRISPR corrected control cell line. The gene expression counts were extracted from the paper. Low expressed genes were removed using filterByExpr() function in the limma package (Ritchie et al., 2015). Gene expression counts were then normalized using the trend approach in the limma package. Genes were next examined for the differential expression across neural progenitor differentiation timepoints (i.e., 4 weeks and 9 weeks) using the limma package with moderated F-statistics.

### Analysis of single cell data

We analyzed the expression patterns of broadly-expressed and brain-specific rASD genes in different neuronal cell types based on available single cell data from two different datasets capturing the gene expression data across brain cell types at prenatal and postnatal ages.

#### Prenatal brain single cell study 1 (Nowakowski et al., 2017)

Processed gene expression data on 4261 single cells from human prenatal brain and associated cell type label of each cell were retrieved (Nowakowski et al., 2017). This single cell dataset provides a relatively higher sequencing coverage compared to other available single cell datasets on human prenatal brain, allowing to better measure the activity of different gene sets in different cell types. Gene expression data were log2(x+1) normalized. For gene set level analysis of broadly-expressed and brain-specific rASD genes, in each sample, we sorted the genes based on their expression level from highest to lowest expression. We next tested separately if the broadly-expressed and brain-specific rASD genes are significantly skewed towards strongly expressed or low expressed genes in each sample using the Wilcoxon-Mann-Whitney test. The *P* values were next z-transformed. The extent of over and under expression in each cell type (i.e., effect size) was measured using the Cohen’s D metric. We used regression models to assess if the effect sizes systematically change across neuronal cell types (i.e., progenitor cells, excitatory and inhibitory neurons) and laminae (i.e., ventricular zone, subventricular zone, and cortical plate). We used the results from this dataset to illustrate the prenatal gene expression patterns of rASD genes in Fig 1G.

#### Prenatal brain single cell study 2 (Li et al., 2018)

Normalized gene expression data on 726 single cells were retrieved. Gene expression data were log2(x+1) normalized. The expression pattern of broadly-expressed and brain-specific rASD genes across neuronal cells were examined by GSEA using Wilcoxon-Mann-Whitney test.

#### Adulthood single cell brain gene expression data (Li et al., 2018)

Unique molecular identifiers (UMI) counts were retrieved (Li et al., 2018). Gene expression data were log2(x+1) normalized. The expression pattern of broadly-expressed and brain-specific rASD genes across neuronal cells were examined by GSEA using a Wilcoxon-Mann-Whitney test.

### Enrichment of rASD genes for the regulators of PI3K/AKT, RAS/ERK, and WNT/ β-catenin signaling pathways

We retrieved the results from a CRISPR knock out library study that investigated the effect of the single gene knockouts on the phosphorylation status of AKT, β-catenin, and ERK 1/2 proteins (Brockmann et al., 2017). Background included Broadly-expressed and brain-specific genes that were included in the study (Brockmann et al., 2017) and were expressed in the developing brain based on the neocortex RNA-Seq data from BrainSpan (BrainSpan, 2016; Kang et al., 2011).

### Analysis of Microglia VPA gene expression data

RNA-Seq data were downloaded from NCBI GEO (GSE109329) (Cho et al., 2019). Expression values were log2(x+1) transformed and quantile normalized. For Entrez genes that were represented by multiple ensemble gene IDs, we retained the gene with highest mean expression level. DE analysis between VPA treatment and controls was performed by limma and moderated t-tests. Genes with fold change >1.5 and Benjamini-Hochberg corrected *P* <0.1 were deemed as DE. To assess the overlap of VPA response in neuronal and microglia, we compared the DE genes in microglia with the reported DE genes in neurons in response to VPA treatment (Chanda et al., 2019) (Fig 4F). To identify the group level response of broadly-expressed and brain-specific rASD genes, we conducted GSEA in each transcriptomic sample using the Wilcoxon-Mann-Whitney test. *P* values were next z-transformed. For each rASD group, we next compared the z-scores between the VPA treated and control samples.

### Expression of broadly-expressed and brain-specific rASD genes in Microglia transcriptome datasets

We examined the expression pattern of broadly-expressed and brain-specific rASD genes in different brain cell types and at different time periods using three available transcriptome datasets:

#### Study 1 (Matcovitch-Natan et al., 2016)

RNA-Seq data from mouse yolk sac progenitors and microglia were retrieved from NCBI GEO (GSE79812). Genes with a TPM normalized expression level >3 were considered as expressed in each sample. The mouse gene IDs were mapped to human orthologs using the Mouse Genome Informatics (MGI) database.

#### Study 2 (Galatro et al., 2017)

RNA-Seq data from brain and microglia isolated from parietal cortex of 39 human subjects were retrieved from NCBI GEO (GSE99074). Genes with a TPM normalized expression level >3 were considered as expressed in each sample.

#### Study 3 (Zhang et al., 2014)

RNA-Seq data on microglia, neurons, astrocytes, oligodendrocytes, and endothelial cells were retrieved from the study. Genes with a TPM normalized expression level >3 were considered as expressed in each sample.

A gene was deemed as expressed in a cell type from each dataset if it was expressed in >50% of the samples related to the cell type in the dataset.

### Expression response of rASD genes to MeCP2, FMR1, and PTEN knockdown

Microarray gene expression data were downloaded from NCBI GEO (GSE47150) (Lanz et al., 2013). Genes with probe intensities below 15 in more than half of the samples were marked as not expressed and were subsequently removed (Lanz et al., 2013). Data was next log transformed and quantile normalized. The mouse gene IDs were mapped to human orthologs using the Mouse Genome Informatics (MGI) database. For human genes with multiple probes in the microarray dataset, we retained the probe with highest mean expression across the dataset. Limma with moderated t-statistics (Ritchie et al., 2015) was used to identify genes dysregulated in the knockdown of MeCP2, FMR1, and PTEN compared to anti-luciferase shRNA treated control cells, separately. Genes with Benjamini-Hochberg corrected *P* <0.1 and fold change >1.5 were deemed as significant in each knockdown background.

### BAF170 knockout gene expression data

Log transformed microarray data were downloaded from NCBI GEO (GSE45629) (Tuoc et al., 2013). Probes with the detection *P* of greater than 0.05 in more than half of the samples were marked as not expressed and were removed. Expression data was next quantile normalized. The mouse gene IDs were mapped to human orthologs using the Mouse Genome Informatics (MGI) database. For human genes that mapped to multiple probes, only the probe with highest mean expression was retained. DE genes in response to knockout of BAF170 were identified using the limma package (Ritchie et al., 2015) with moderated t-statistics. Genes with the fold change >1.3 and the Benjamini-Hochberg corrected *P* <0.1 were deemed as DE (414 DE genes).

### SOX5 over-expression gene expression data

RNA-Seq read counts were downloaded from the NCBI GEO with the accession GSE89057 (Parikshak et al., 2016). Low expressed genes were removed using the filterByExpr() function in the limma package (Ritchie et al., 2015). Read counts were next normalized and DE genes were identified using the trend approach coupled with the moderated t-statistics implemented in the limma package. Genes with the fold change >1.5 and Benjamini-Hochberg *P* <0.05 were deemed as significant. The original paper on the SOX5 over-expression transcriptome data had included the first two principal components (PCs) of transcriptome data as additional co-variates in the DE analysis pipeline. We tested the inclusion of the PCs in the regression model of the limma and observed highly concordant results with the model that considers only the genotype (i.e., over-expression of SOX5 and normal controls). Therefore, since the covariates that the two PCs were measuring were ambiguous in nature, we elected not to include the two PCs in our DE analysis.

### ASD postmortem brain gene expression data

DE analysis results comparing the expression level of 16,398 genes between ASD and control cortex were downloaded (Parikshak et al., 2016). Gene IDs were converted from Ensembl gene IDs to Entrez IDs using the BioMart database. Genes with Benjamini-Hochberg corrected *P* <0.1 were deemed as DE.

### Hematopoietic differentiation gene expression data

Normalized microarray data were downloaded from GSE24759 (Novershtern et al., 2011). In this dataset, the gene expression data were already corrected for batch effects using Combat. The expression of each gene was further standardized to have mean of zero and a variance equal to one. To examine the conservation of the neurodevelopmental network of broadly-expressed and brain-specific rASD genes, we defined the interaction preservation score as: 

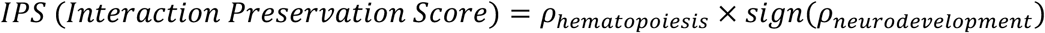

Where 

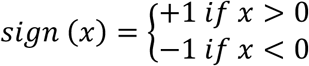

and *ρ*_*hematopoiesis*_ represents the Pearson’s correlation coefficient of the interaction during the hematopoiesis and *ρ*_*neurodevelopment*_ represents the Pearson’s correlation coefficient of the interaction during the prenatal and early postnatal time periods (8pcw to 1yr). We calculated the *IPS* score for each interaction in the neurodevelopmental network of broadly-expressed and brain-specific rASD genes. As background, we constructed a global neurodevelopmental gene network using the same approach as the rASD genes. We next compared the *IPS* scores from the rASD networks with that of the global neurodevelopmental network as the background. Since, by design, broadly-expressed rASD genes are strongly expressed in other tissues including the blood, we limited all three networks to those genes that are expressed during the hematopoiesis differentiation as well, to maintain the comparability.

For GSEA, in each sample, we sorted the broadly-expressed and brain-specific rASD genes based on their normalized expression from the highest to the lowest, separately. We next examined if the XP-broad or XP-specific genes are randomly distributed in the ranked list or their distribution is skewed towards the genes with high or low relative expressions as judged by the Wilcoxon-Man-Whitney rank sum test. To examine the relationship between the expression patterns of XP-broad and XP-specific genes, we first standardized the expression of each gene to have a mean of zero and standard deviation of one. We next performed GSEA and z-transformed the *P* and examined the Pearson’s correlation coefficient between the z-scores of XP-broad and XP-specific genes across the samples.

