## Supplementary Figures for "Autism genetics perturb prenatal neurodevelopment through a hierarchy of broadly-expressed and brain-specific genes"

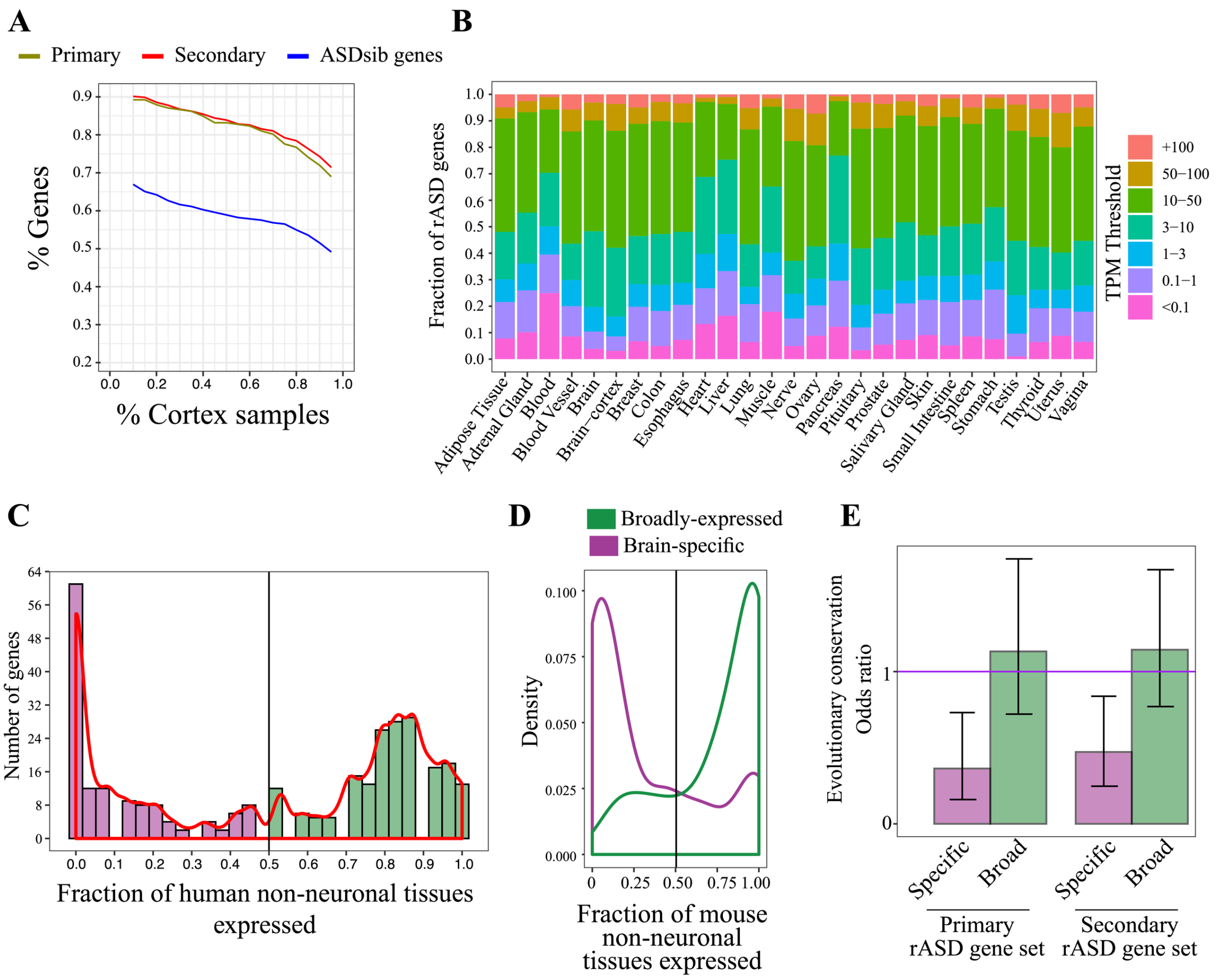


**Fig S1. ASD risk genes are composed of both broadly-expressed and brain-specific genes.**
To assess the robustness of the results, we repeated our analyses on a secondary rASD gene set, defined based on a different criteria (Methods). All figures illustrated here demonstrate the analysis results on this secondary rASD gene set, except for panels A and E that includes results on both the primary and the secondary rASD gene sets. **A)** rASD genes show stronger expression in the adult cortex compared to the genes identified as mutated in the typically developing siblings of ASD individuals (ASDsib genes). The X-axis represents the fraction of the adult cortex samples, and the Y-axis represents the fraction of genes that show expression in a given fraction of cortex samples. **B)** A group of rASD genes are strongly expressed across tissues. The figure represents the expression pattern of rASD genes across human tissues based on the GTEx dataset. The expression of rASD genes were categorized based on their TPM normalized expression levels (Methods). **C)** rASD genes demonstrate a bimodal expression pattern with some broadly-expressed across tissues, while others showing brain-specific gene expression patterns. For the rASD genes that were expressed in the adult cortex, we calculated the fraction of non-neuronal tissues in which the genes show expression. The higher values indicate that the genes are more broadly-expressed across human tissues. **D)** Gene expression patterns of rASD genes are conserved in mouse tissues. The expression patterns of broadly-expressed and brain-specific rASD gene homologs were examined across mouse tissues. The X-axis demonstrate the fraction of non-brain tissues in which the homologs show expression. As illustrated, homologs broadly-expressed rASD genes tend to be broadly-expressed across mouse tissues as well. The pattern reverses for the brain-specific rASD genes. **E)** Brain-specific rASD genes are not evolutionary conserved in yeast. The broadly-expressed and the brain-specific rASD genes were defined based on their gene expression patterns across human tissues. To examine the functional relevance of this categorization, the evolutionary conservation of the broadly-expressed and brain-specific genes were examined in *Saccharomyces cerevisiae*. The rational was that since *S. cerevisiae* do not have neurons, the genes that are specific to the brain functioning are expected to be depleted in this organism. Gene ortholog mappings were retrieved from the BioMart database. Additional analyses further supported that the brain-specific genes are highly enriched for the genes that are specific to neurons (Fig S9).

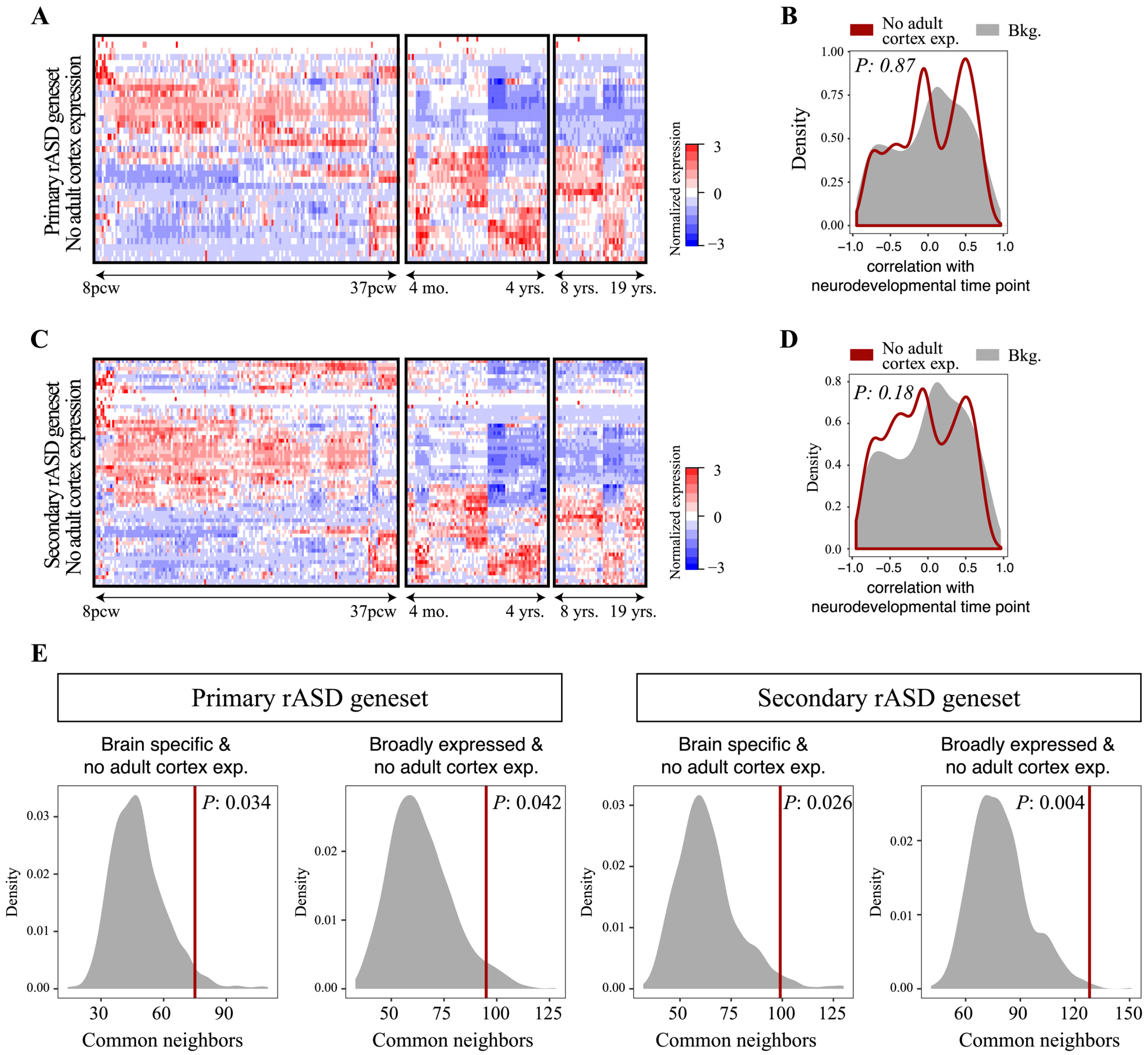


**Fig S2. ASD risk genes not expressed in the adult cortex are associated similarly with both broadly-expressed and brain-specific rASD genes.**
Broadly-expressed and brain-specific rASD genes demonstrated contrasting gene expression patterns during the prenatal brain development. Specifically, broadly-expressed genes were enriched among the early expressing genes with a peak expression at the early neurodevelopmental times points, while most of the brain-specific genes exhibit peak expression at the later neurodevelopmental states. Moreover, broadly-expressed and brain-specific rASD genes were involved in dense subnetworks that were separate from each other. We leveraged these two sets of information to examine if the genes not expressed in the adult brain cortex show bias towards brain-specific or broadly-expressed genes. **A-D)** Genes not expressed in the adult cortex do not show strong enrichment towards either early or late expressing genes. The panels A and B are based on the analysis of primary rASD gene set, and the panels C and D are based on the secondary rASD gene set. The correlation of gene expression patterns with the prenatal and early postnatal time periods (8 pcw to 2 years after) were calculated using the biweight midcorrelation efficiently metric. **E)** rASD genes not expressed in the adult cortex are connected in a similar extent to both networks of broadly-expressed and brain-specific rASD genes.

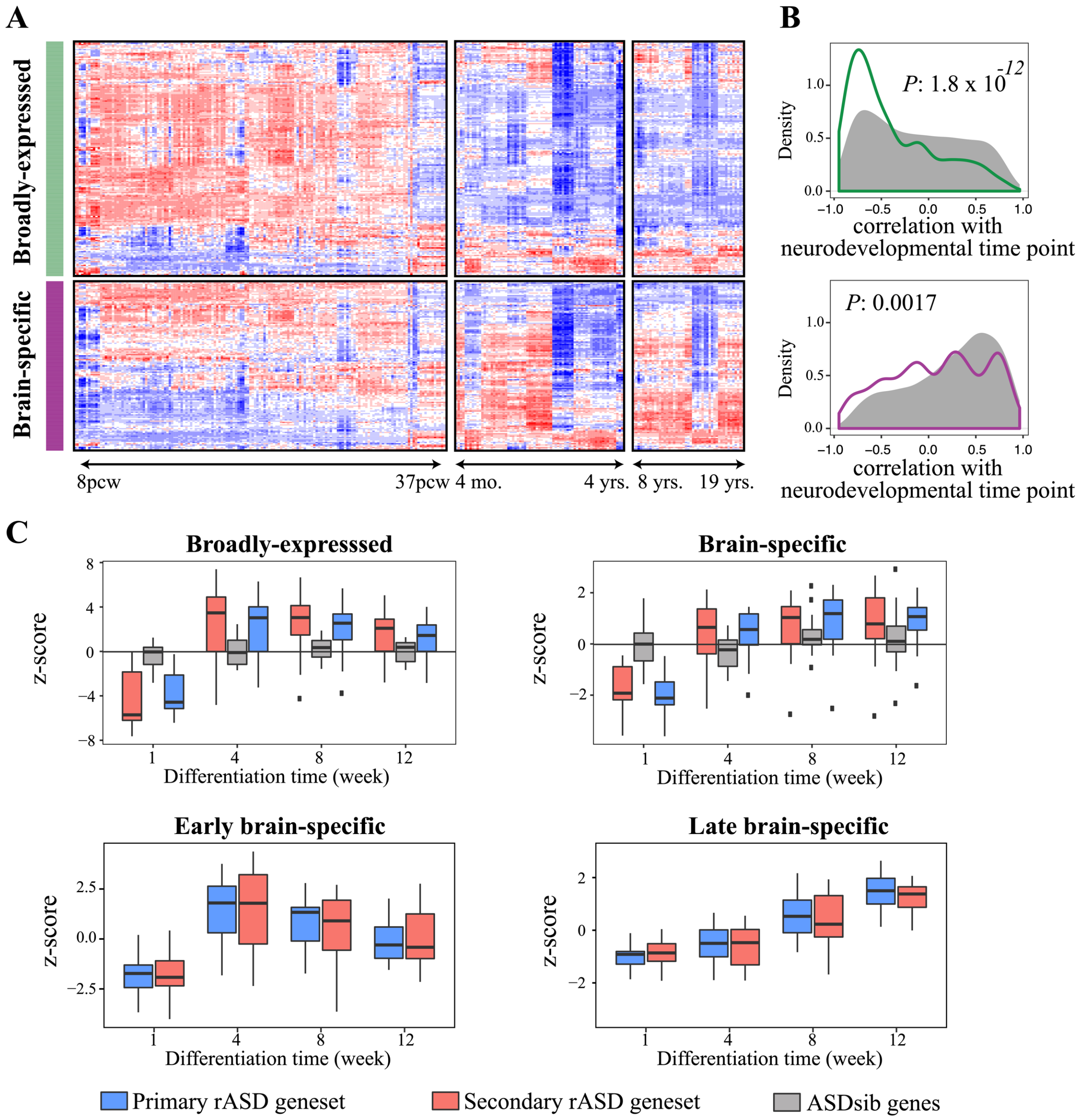


**Fig S3. Broadly-expressed and brain-specific ASD risk genes are part of different transcriptional programs.**
**A)** Broadly-expressed and brain-specific rASD genes demonstrate different expression patterns during the neurodevelopment. The analysis is based on the secondary rASD gene set. The figures illustrate the expression pattern of rASD genes from 8 pcw to 19 years old. The expression of each gene was standardized to have mean of zero and standard deviation of one. **B)** The comparison of temporal expression pattern of broadly-expressed and brain-specific rASD genes with the background composed of all broadly-expressed and brain expression genes. As illustrated the rASD gene groups show significantly different expression patterns than the background. The correlation of each gene with the neocortex temporal developmental stages during prenatal and early postnatal ages (8 pcw to 2 years) was calculated using the biweight midcorrelation metric and considering the time points as ordinal variables. Broadly-expressed and brain-specific ASDsib genes did not show significantly different neurodevelopmental expression patterns compared to the corresponding background genes ($P > 0.05$; data not shown) **C)** Gene expression patterns of broadly-expressed and brain-specific rASD genes during in vitro neuronal differentiation. As illustrated, the expression patterns of gene groups is similar to their patterns during the fetal brain development, suggesting the relevance of the prenatal rASD gene expression patterns in the brain to the neural differentiation processes. GSEA in each sample was conducted by sorting all broadly-expressed and brain-specific genes based on their expression levels, separately. We next assessed whether broadly-expressed and brain-specific rASD genes show significant bias towards high expressing or low expressing genes in the corresponding gene expression sets. The figure illustrates the z-scores from the Wilcoxon-Mann-Whitney test.

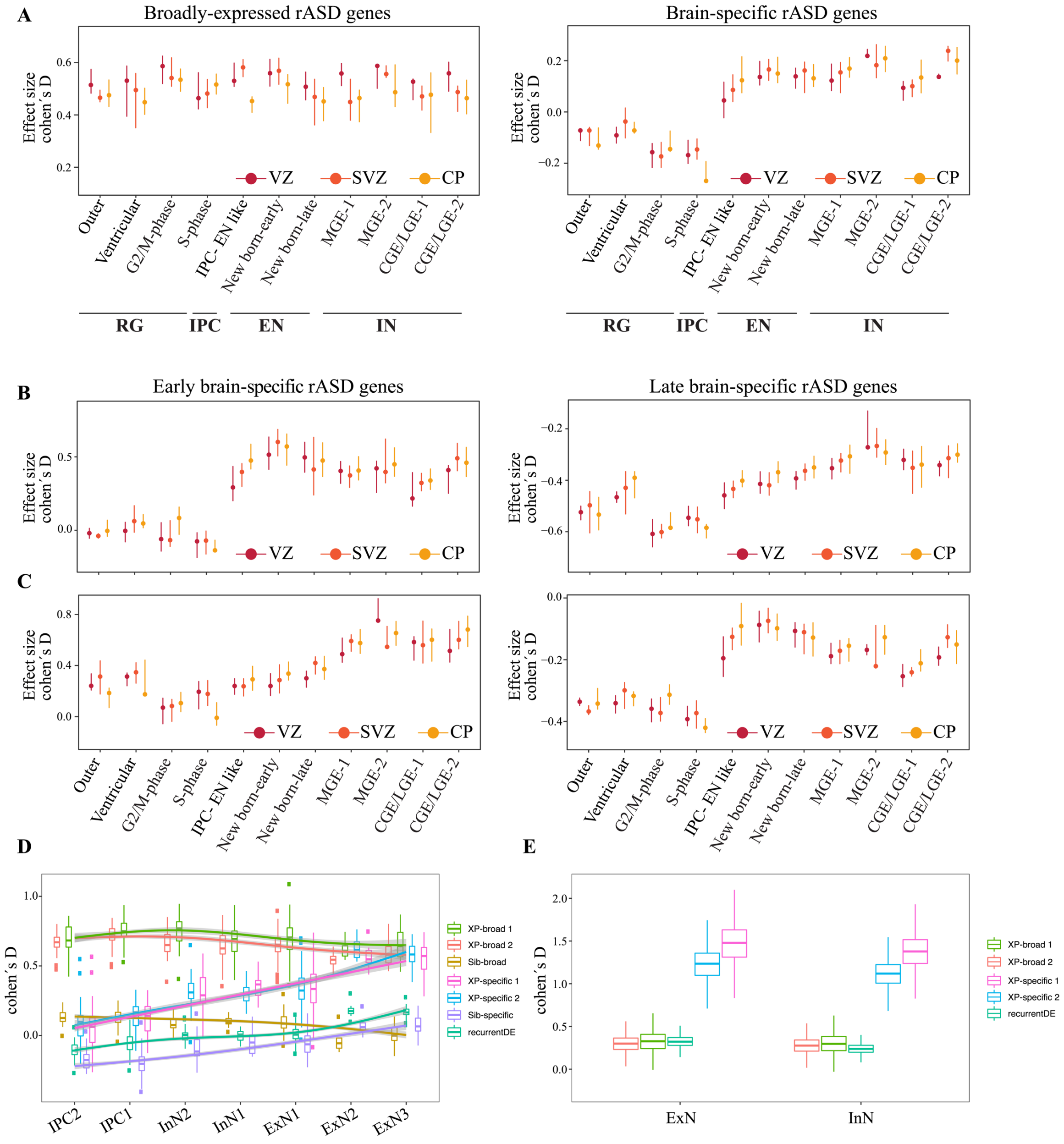


**Fig S4. Single cell gene expression data reveals the differences in the gene expression programs of broadly-expressed and brain-specific ASD risk genes.**

**A)** Gene expression of broadly-expressed rASD genes are significantly reduced during the transitions from ventricular zones (VZ) to subventricular (SVZ) and cortical plate (CP) regions. However, the expression of the brain-specific genes are mostly increased due to cell differentiations from proliferative to neuronal cells. The analysis is based on the secondary rASD gene set. GSEA with a follow up effect size estimation was conducted on the expression levels from each cell to identify the expression pattern of broadly-expressed and brain-specific rASD genes. The expressed genes in each cell were considered as the background. Circles represent the median effect size, and the lines illustrate the associated interquartile range. **B)** Early and late brain-specific rASD genes show similar expression trends during the neuronal differentiation. However, the early expressing genes show stronger base expression levels during the 10 to 20 pcw. The analysis is based on the primary rASD gene set. To be able to compare between the expression levels of gene sets composed of different number of genes, we calculated Cohen's D effect sizes. **C)** Similar to panel B, but on the secondary rASD gene set. **D)** Prenatal gene expression patterns of broadly-expressed and brain-specific rASD genes based on a secondary single cell dataset (Li et al., 2018). As illustrated, the expression of the brain-specific rASD genes are strongly regulated by cell differentiation events. **E)** Brain-specific rASD genes are more strongly expressed in the adult neurons compared to the broadly-expressed rASD genes. The single cell data on adult cortex was retrieved from (Li et al., 2018). XP-broad 1 and XP-specific 1: Primary rASD gene set. XP-broad 2 and XP-specific 2: Secondary rASD gene set. ExN: Excitatory neuron; InN: Inhibitory neuron.

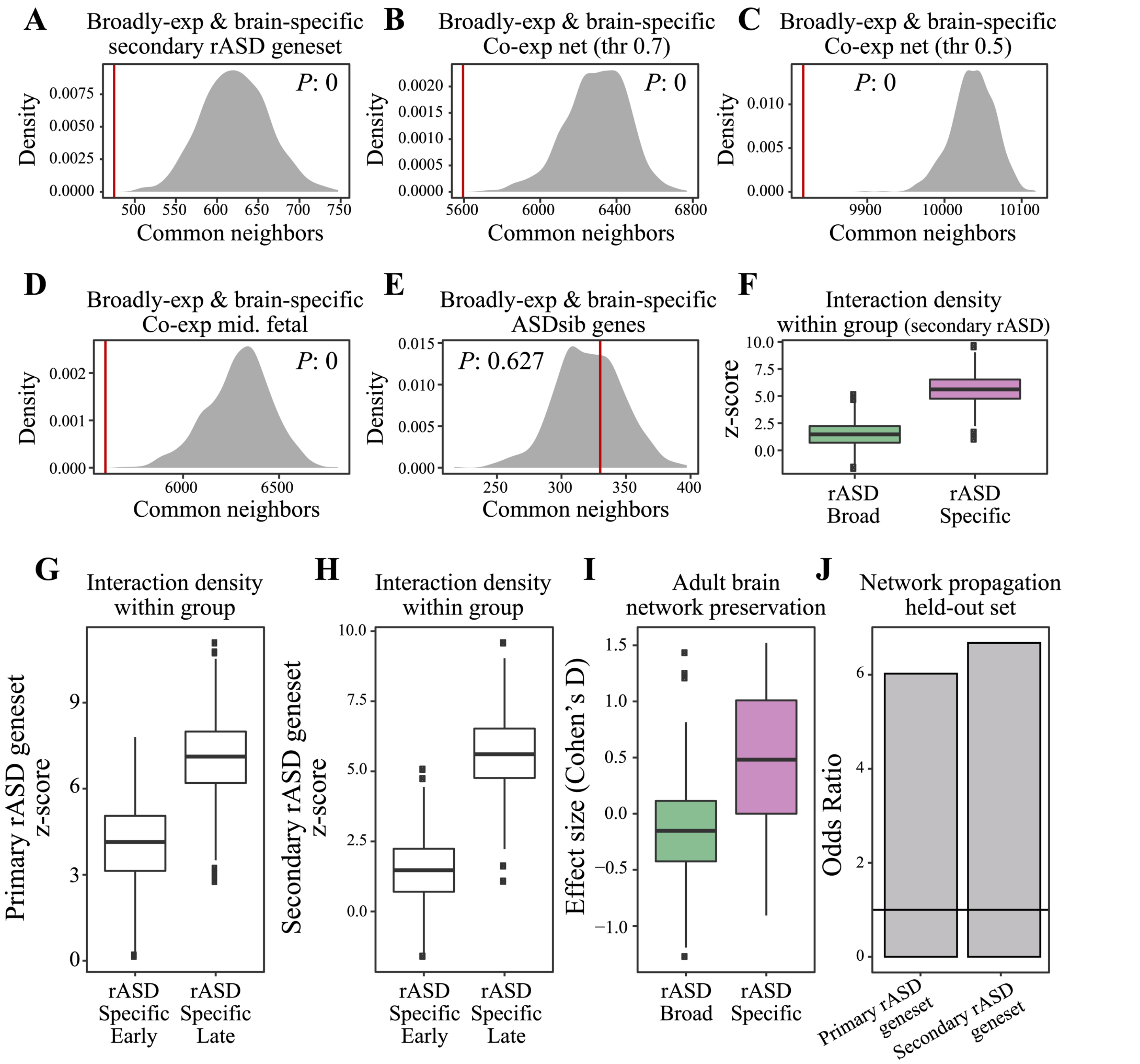


**Fig S5. Broadly-expressed and brain-specific ASD risk genes are involved in separate gene networks.**
**A)** Gene networks of broadly-expressed and brain-specific rASD genes are depleted for common gene neighbors. This analysis is based on the secondary rASD gene set. The network is based on high confidence regulatory and physical interactions that show strong co-expression during the prenatal and early postnatal brain development (Methods). The number of common gene neighbors between the networks of broadly-expressed and brain-specific rASD genes (red line) were next compared to a background distribution to calculate the empirical *P*. The background distribution was constructed by identifying the number of common gene neighbors between randomly selected broadly-expressed and brain-specific genes with the node degree distribution as broadly-expressed and brain-specific rASD genes, respectively. **B)** Analysis approach similar to panel A, but the neurodevelopmental network was constructed purely based on the co-expression of the genes during prenatal and early postnatal ages (All interactions with co-expression strength of >0.7 were included in the analysis; unsigned Pearson's correlation). This analysis indicates that the separation of the broadly-expressed and brain-specific genes is not due to existence of potential missing knowledge on the interactions between the broadly-expressed and the brain-specific genes. The analysis is based on the primary rASD gene set. **C)** Similar to panel B, but the neurodevelopmental network was constructed by considering interactions with co-expression strength of >0.5 (unsigned Pearson's correlation). The analysis is based on the primary rASD gene set. **D)** Similar to panel B, but the co-expression network was constructed by considering the neocortex gene expression data during 12 to 24 pcw. **E)** In contrast to the rASD genes, the neurodevelopmental networks of broadly-expressed and brain-specific ASDsib genes are not significantly separate. **F)** brain-specific and broadly-expressed rASD genes are involved in dense gene networks. The analysis is based on the secondary gene set. **G-H)** The

inter-connectivity of late brain-specific rASD genes is stronger than the early brain-specific rASD genes. The analysis is based on the primary and secondary rASD gene sets, respectively. **I)** Neurodevelopmental network of brain-specific genes is significantly conserved in the adult cortex. The preservation of neurodevelopmental networks of rASD genes were assessed using the normalized expression data from GTEx consortium. A gene-centric approach was employed to examine the preservation of the gene networks by comparing the co-expression strength of interactions of each rASD gene in the adult cortex with a background distribution using the Kolmogorov-Smirnov test. Analysis is based on the secondary rASD gene set. **J)** The employed network propagation approach successfully identifies the held-out rASD genes. Gene labels of 20% or rASD genes were randomly set to unknown prior to the application of network propagation. Next, we examined the enrichment of the held-out genes among the genes that are identified as significantly connected to rASD genes through the network propagation.

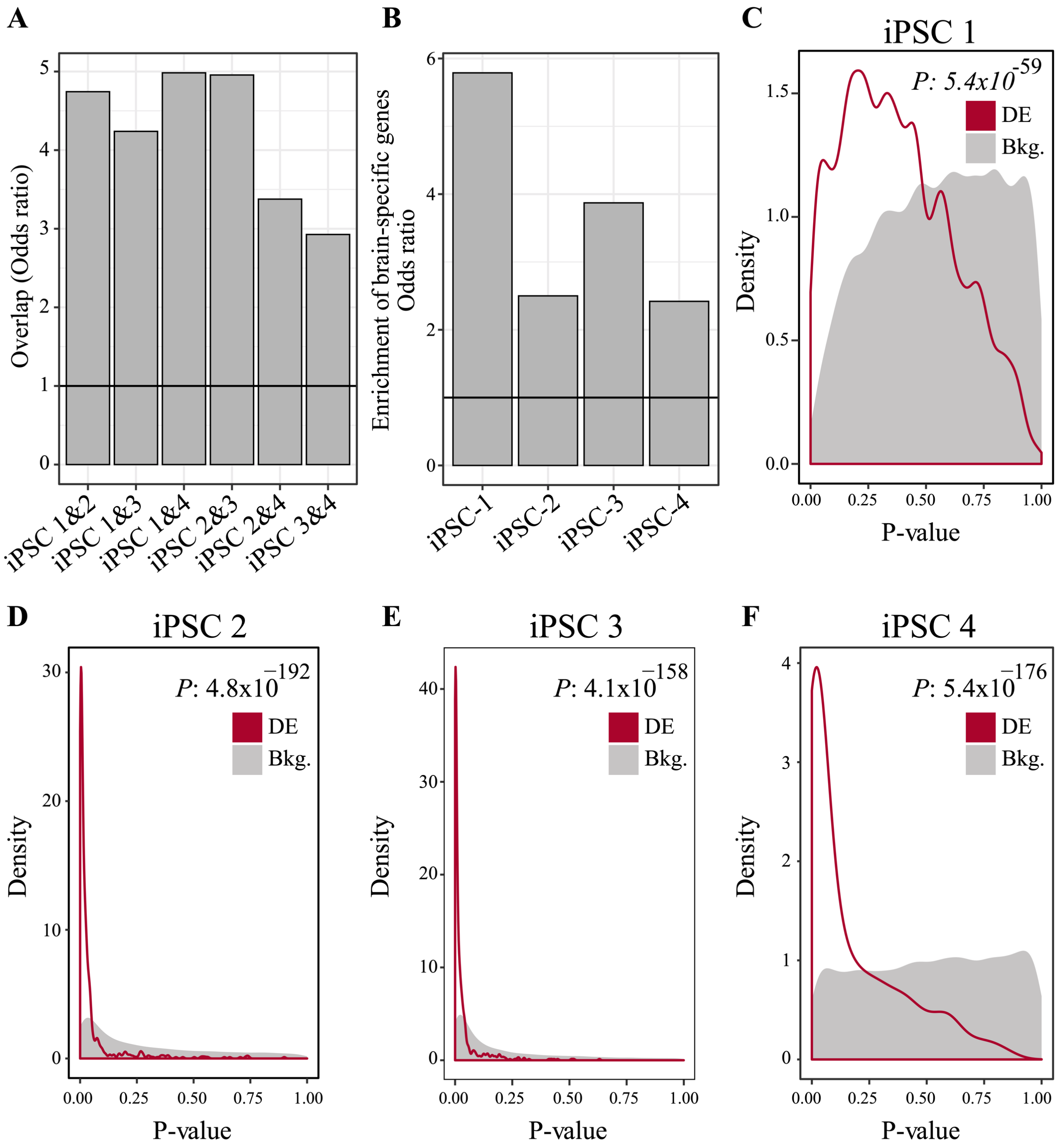


**Fig S6. Common genes are perturbed in neuronal models of ASD**

**A)** Pairwise overlap analysis of DE genes from four independent hiPSC studies on ASD. DE genes from each study include the genes that are perturbed during the differentiation of neural progenitors to neurons. All comparisons have FDR <0.1 (Fisher's exact test). **B)** Genes perturbed in hiPSC-derived neuronal models of ASD are enriched for the brain-specific genes. **C-F)** The identified 599 recurrent DE genes show strong perturbations in each of the four independent transcriptome datasets on hiPSC-derived neuronal models of ASD. The X-axis indicates the *P* that the genes are DE in hiPSC-derived neural progenitor and neurons from individuals with ASD. The transcriptome data in panel C is from an idiopathic ASD cohort. Transcriptome data in panels D and E relate to ASD cohorts with macrocephaly. The transcriptome data in panel F include data from an ASD individual with a SHANK2 loss-of-function mutation. See methods for the details on the data analysis. iPSC 1: (Liu et al., 2017); iPSC 2: (Mariani et al., 2015); iPSC 3: (Schafer et al., 2019); iPSC 4: (Zaslavsky et al., 2019).

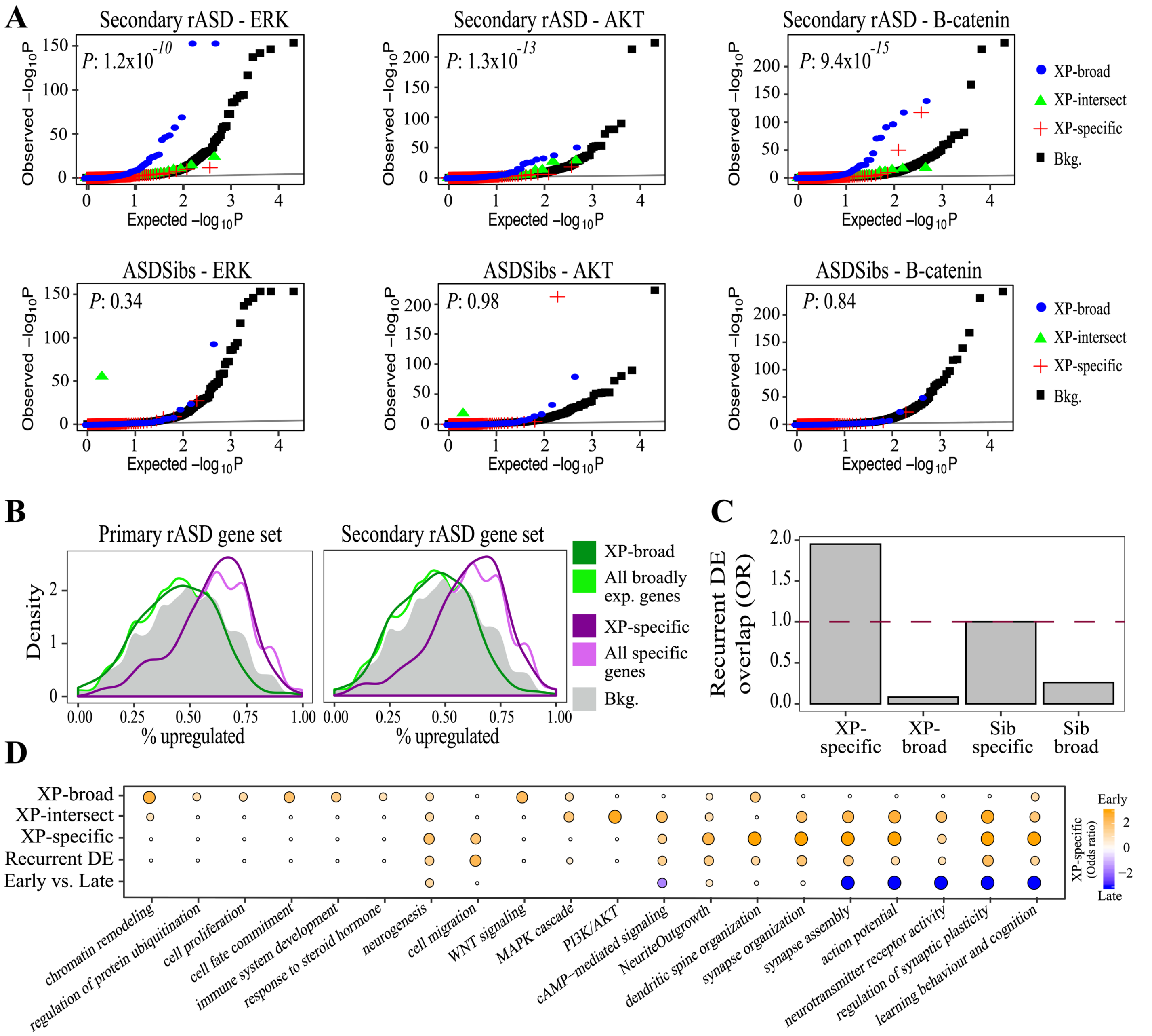


**Fig S7. Neurodevelopmental gene networks of broadly-expressed and brain-specific rASD genes are connected with the recurrent DE genes in different ways.**

**A)** XP-broad network is significantly enriched for the regulators of PI3K/AKT, RAS/ERK, and WNT/$\beta$-catenin signaling pathways. The results are based on the secondary rASD gene set and ASDsib genes. **B)** XP-specific and brain-specific genes in general, show an upregulation trend in hiPSC-derived neuronal models of ASD. The left and right panels are based on the primary and the secondary rASD gene sets, respectively. **C)** XP-specific network significantly overlaps with the recurrent DE genes from hiPSC-derived neuronal models of ASD. The analysis is based on the secondary rASD gene set. **D)** XP-broad network is significantly enriched for the processes involved in the initial stages of neural development, while the XP-specific network exhibits enrichment for the processes related to the neural maturation, synaptogenesis, and synapse functioning. Note that the recurrent DE genes show similar enrichment patterns with the XP-specific genes. Moreover, XP-intersect genes that connect the XP-broad network to the XP-specific network are strongly enriched for the signaling pathways. The early/late category represent the relative enrichment of the processes between the early and late expressing rASD genes.

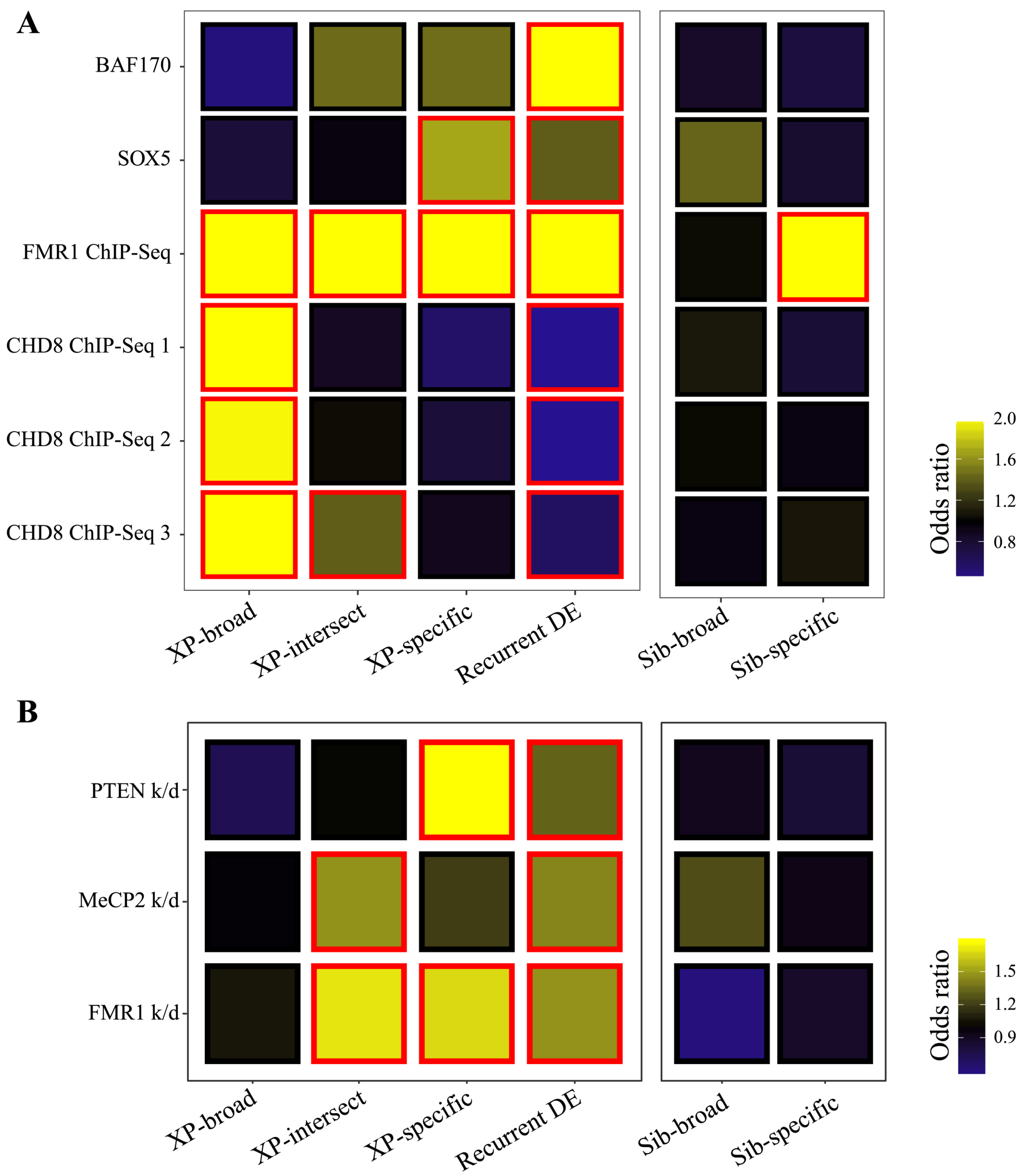


**Fig S8. PI3K/AKT, RAS/ERK, and WNT/$\beta$-catenin signaling pathways modulate the expression of XP-specific and recurrent DE genes.**

**A)** Effect of genetic aberration of high confidence rASD genes on the neurodevelopmental gene networks of broadly-expressed and brain-specific genes. The colors illustrate the odds ratio, and the red borders demonstrate significant odds ratios (FDR <0.1). ChIP-Seq data on the CHD8 were retrieved from three studies CHD8 ChIP-Seq 1: (Gompers et al., 2017); CHD8 ChIP-Seq 2: (Sugathan et al., 2014), and CHD8 ChIP-Seq 3: (Cotney et al., 2015). ChIP-Seq regulatory targets of FMR1 were retrieved from (Darnell et al., 2011). The genes perturbed in response to the BAF170 knockout and SOX5 overexpression were identified by analysis of available transcriptome data from (Tuoc et al., 2013) and (Parikshak et al., 2016), respectively. The results are based on the secondary rASD gene set and ASDsib genes. **B)** PI3K/AKT, RAS/ERK, and WNT/$\beta$-catenin signaling pathways modulate the expression of XP-specific and recurrent DE genes. The three genes of PTEN, FMR1, and MeCP2 modulate the three signaling pathways. Genes perturbed in mouse neurons in response to the shRNA-induced knockdown of PTEN, FMR1, and MeCP2 were identified by the analysis of available microarray gene expression data (Lanz et al., 2013). Analyses are based on the secondary rASD gene set and ASDsib genes.

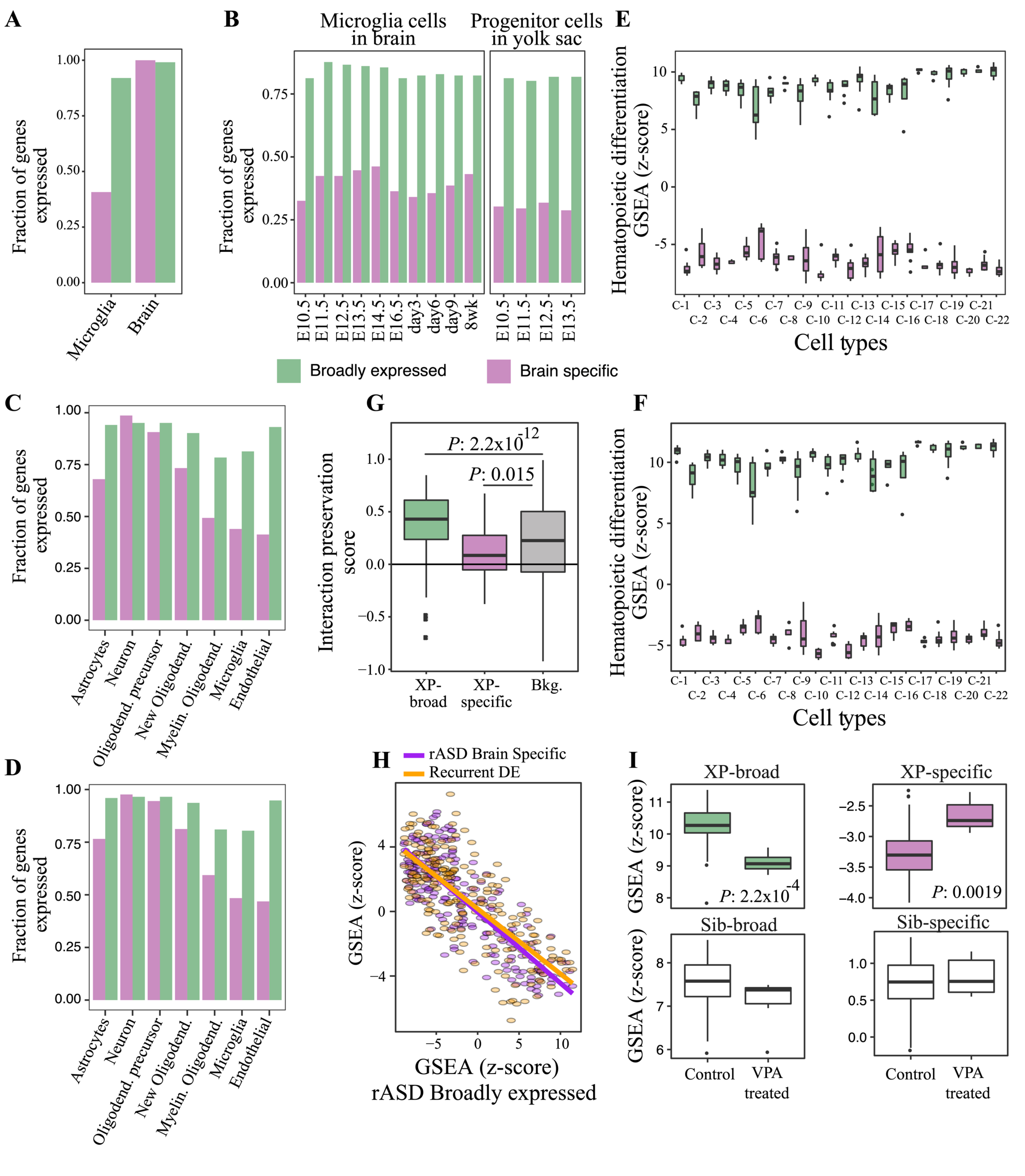


**Fig S9. Broadly-expressed rASD genes are strongly expressed in non-neuronal brain cells.**
**A)** Expression level of broadly-expressed and brain-specific rASD genes in brain and microglia isolated from parietal cortex of 39 human subjects *(Galatro et al., 2017)*. Data is based on the primary rASD gene set. **B)** Expression of broadly-expressed and brain-specific rASD genes in mouse yolk sac progenitors and microglia *(Matcovitch-Natan et al., 2016)*. Analysis is based on the secondary rASD gene set. **C-D)** Expression of broadly-expressed and brain-specific rASD genes in microglia, neurons, astrocytes, oligodendrocytes, and endothelial cells *(Zhang et al., 2014)*. Analysis is based on the primary (panel C) and secondary (panel D) rASD gene set. **E-F)** Broadly-expressed rASD genes are strongly expressed during the hematopoiesis (Novershtern et al., 2011). The

Y-axis represent the z-score from the GSEA of the two rASD gene groups using the Wilcoxon-Mann-Whitney test. The analysis is based on the primary (panel E) and secondary (panel F) rASD gene sets. **G)** Neurodevelopmental network of broadly-expressed rASD genes is conserved during the hematopoiesis differentiation. Interaction preservation score (IPS) measures the strength and preservation of the neurodevelopmental interactions in during the hematopoiesis differentiation (see Methods). The analysis is based on the secondary rASD gene set. **H)** Broadly-expressed genes exhibit an anti-correlated expression pattern with the brain-specific and recurrent DE genes during the hematopoiesis differentiation. GSEA was conducted on the normalized gene expression levels where the expression of each gene was standardized to have a mean of zero and standard deviation of one. The *P* from the Wilcoxon-Mann-Whitney test were next z-transformed. **I)** Broadly-expressed and brain-specific rASD genes show opposite responses to the VPA treatment in the microglia. In each sample, the group level expression levels were calculated by GSEA using Wilcoxon-Mann-Whitney test. The analysis is based on the secondary rASD gene set and the ASDsib genes.

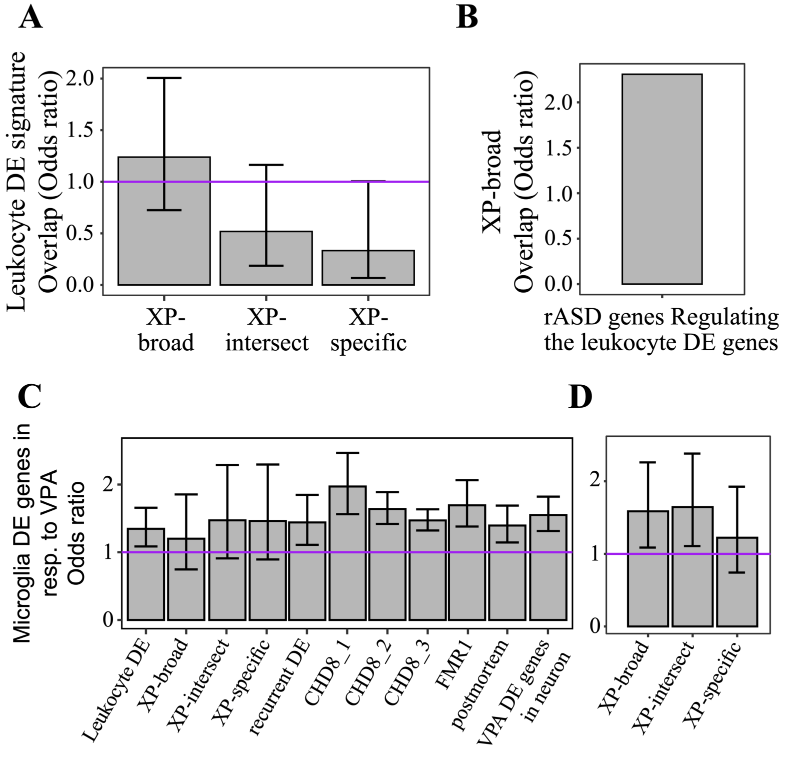


**Fig S10. Genes perturbed in microglia in response to VPA overlap with the genes perturbed in leukocytes of individuals with ASD from the general population.**

**A)** The leukocyte DE signature does not overlap with the XP-specific and XP-broad genes. **B)** The regulators of the leukocyte DE signature overlap with the broadly-expressed rASD genes. The enrichment for the broadly genes were assessed with Fisher's exact test. **C-D)** The Figure demonstrates the overlap of DE genes in response to VPA treatment in microglia with the gene sets implicated in ASD. As illustrated, the VPA-induced DE genes in microglia show an overlap with the DE signature from leukocytes of idiopathic individuals with ASD from general population, suggesting an overlap in the perturbed mechanisms between the two sets. The analysis is based on the secondary rASD gene set. The plots demonstrate odds ratios and 95% confidence intervals. The analysis is based on the primary (C) and secondary (D) rASD gene sets.
